# Allosteric signal initiation and communication in neuromuscular acetylcholine receptors

**DOI:** 10.64898/2026.08.02.742316

**Authors:** Pradeepti Kampani, Tapan K. Nayak

## Abstract

Acetylcholine receptors (AChRs) expressed at the nerve-muscle synapses are prototypic allosteric receptors that shuttle between a resting **C**losed (C) and active **O**pen (O) states. ACh binding at the neurotransmitter binding sites (TBS) leads to opening of the ‘gate’ in the channel pore that regulates ion flow. How agonist binding to the TBS communicates the allosteric signal to the channel gate located ∼50 Å away is not understood. In the absence of agonists, the wild-type and mutant AChRs, including those causing congenital myasthenia syndrome, show constitutive gating. However, whether allosteric activation pathways in the presence vs absence of agonists are identical, is debatable. Here, by using a combination of kinetic modelling, phi (ϕ)- and activation energy (ΔΔG^ǂ^) estimation for >60 residues from single channel current recordings and molecular dynamics simulations, we show the existence of parallel gating pathways (‘major’ and ‘minor’) in the unliganded AChRs and activation pathways are non-identical for the liganded vs unliganded AChRs. Kinetic analysis of the minor gating and correlation between the state residence probabilities vs C-loop conformations suggest, 1. the C-loop capping triggers allosteric communication independent of the presence of an agonist and, 2. the minor gating is a pre-existing allosteric pathway which is preferentially chosen in the presence of agonists. Further, we show the presence of a continuous ‘live-wire’ like allosteric network in the liganded receptor between the TBS and the gate constituted by residues which lower the activation energy barrier by >-4 kcal/mol. In contrast, in the unliganded AChRs, the major gating seems to initiate from ‘hub’ residues in the allosteric network. The results presented here provide novel insight into the mechanism of allosteric signal initiation and communication in AChRs.

## Introduction

The nicotinic acetylcholine receptors (AChRs) expressed at the neuromuscular junction synapses (NMJs) belong to a superfamily of pentameric ligand gated ion channels (LGICs). They are comprised of five subunits; (α1)_2_βδε (adult)/ γ (fetal) with the neuro-transmitter binding sites (TBS) located at the interfaces between αδ and αε/γ. AChRs are allosteric proteins which shuttle between a non-conducting resting **C**losed and an active **O**pen conformational state (1–4). Agonists bind, respectively, with low and high affinities to C vs O states, thus tilting the equilibrium in the favour of the O state. The local C→O transition at the TBS is followed by a cascade of global conformational change in the receptor, known as ‘gating’, that leads to the opening of the channel pore and ion conduction. In the absence of agonists, the AChRs open rarely, with an infinitesimally small probability (*P_o_*<10^-6^) (a.k.a. constitutive gating). Many mutations that cause congenital myasthenia syndrome (CMS) alter the NMJ current responses to ACh by altering the constitutive channel gating(5–7).

A simple thermodynamic cyclic scheme (2) based on allosteric MWC theory (3) (see Fig. 1a) describes satisfactorily the AChR channel function. In the cycle, the vertical and horizontal arms represent gating and binding, respectively. The steady state kinetic rate constants, equilibrium constants (*E_n_* for gating in the presence of *n* agonist molecules and *K_d_*/*J_d_* = equilibrium dissociation constants for the low and high-affinity states, respectively) and Gibb’s free energy changes (ΔG=-0.59*ln (equilibrium constant) at room temperature) associated with agonist binding and channel gating, have been accurately measured by single-channel patch-clamp experiments (8). But the interaction of an agonist with the TBS and the ensuing ‘signal’ to the rest of the receptor about an agonist occupying the binding pocket are pre-steady state events that happen in nanosecond ns-μs time scale (9, 10). We refer to these set of events at the transition state prior to the onset of the global gating as ‘allosteric communication’. Due to the experimental limitations in studying fast events, allosteric communication studies have been challenging (11, 12).

**Figure 1.**
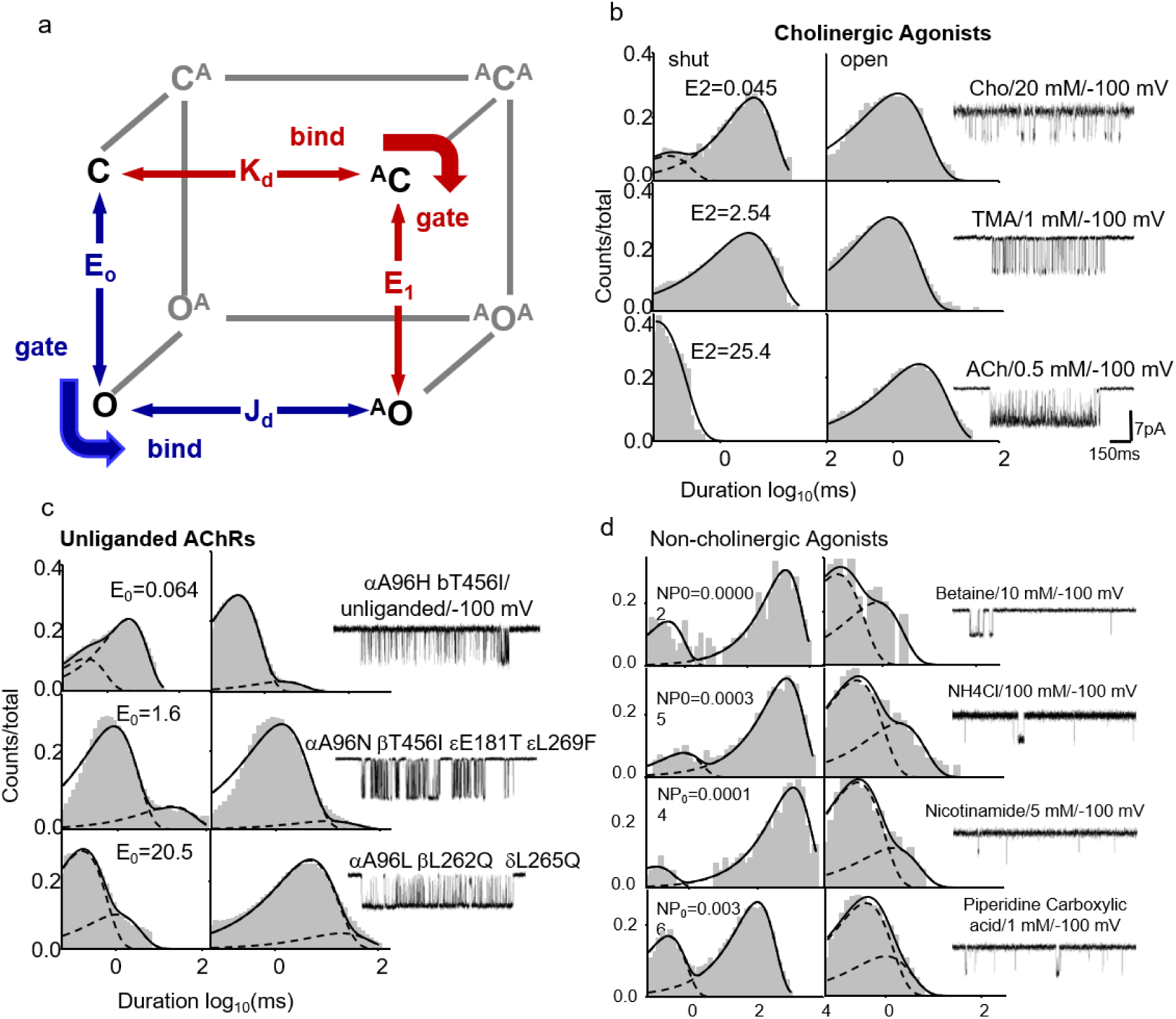
Thermodynamic and kinetic features of agonist-dependent and unliganded AChR gating. **(a)** Thermodynamic cycle. Horizontal and vertical transitions represent agonist binding and receptor gating, respectively. Monoliganded gating is highlighted and shown as the front face of the thermodynamic cube. C and O: Resting closed and active open states, respectively; A: agonist; AC: low-affinity agonist-bound closed state; and AO: high-affinity agonist-bound open state. Bind–gate and gate–bind pathways are shown, together with their associated equilibrium constants. Microscopic reversibility requires that the net free-energy change around a closed thermodynamic cycle is zero. Consequently, ΔG_1_ – ΔG_0_ = ΔG_HA_ − ΔG_LA_ = ΔG_B_, where ΔG₁, ΔG₀, ΔG_HA_, and ΔG_LA_ are the free energy associated with liganded, unliganded, high-affinity- and low-affinity binding. ΔG_B_ = the net agonist energy. **(b)** Effect of cholinergic agonists on AChR gating. Representative single-channel current recordings and the corresponding closed- and open-duration distributions. Solid curves superimposed on the duration histograms represent probability-density functions obtained by fitting the single-channel current data to simple C↔O kinetic models. Note the simple kinetics comprising one closed state and one open state. **(c)** Kinetic complexity of unliganded AChR gating. Representative single-channel current recordings and the corresponding dwell-time distributions showing the kinetic complexity and heterogeneity of unliganded AChR activity. Under each background-mutation condition, multiple C and O states were required to describe the duration distributions. **(d)** Effects of non-cholinergic agonists on the adult-type AChRs. Representative single-channel current recordings and dwell-time distributions obtained in the presence of shown agonists. In each condition, the single-channel activity and duration distributions exhibit complex kinetic behaviour that differs from the relatively simple cholinergic agonist-bound activity shown in panel Fig. 1b and resembles the heterogeneous unliganded activity shown in panel 1c.

Agonist binding to the low affinity C state involves local conformation changes (‘induced fit’) prior to C↔O conformational switch. It is evidenced by: (i) variation of agonist association (*k_on_*) to the C state for different cholinergic agonists (13) (ii) temperature dependence of *k_on_* owing to an enthalpy driven low affinity complex formation (iii) linear correlation between affinity and efficacy for structurally related classes of agonists suggesting that C↔CA and CA↔OA are parts of the same gating continuum in the AChRs (14), GABA receptors (15) and NMDA receptors (16). Structurally, low-affinity binding is associated with distinct conformational changes in the loops that constitute the binding pocket (17). Especially, the C-loop has been shown to undergo a capping movement during agonist binding to the low-affinity state of the receptor (18), while antagonists and toxins fail to elicit such conformational changes (19). On the other hand, agonist association to the high-affinity, constitutively active O state of AChRs has been shown to be diffusion limited over a barrier-less transition state (8) and involves reorientation of the agonist in the TBS (20). Constitutive gating of AChRs is complex with multiple O states (21, 22). It is unclear how an agonist interacts with a conformationally heterogeneous constitutively active O states and if the mechanism of agonist binding to the O vs C states is similar.

To study the mechanism of allosteric communication, it is essential to understand the nature of the transition state of the receptor. The transition state ensemble (TSE) is constituted by innumerable short-lived intermediate states between the C and O stable end states. Some of them, which are relatively more stable, including ‘Flip’ (23) and ‘Prime’ (24) states, have been inferred from experimental single channel data. However, the energetics of the transition state are difficult to investigate. The position of the transition state, however, has been studied in different proteins (25, 26) including AChRs by linear free energy equilibrium relationships (LFERs;(27)) and phi (ϕ) value analysis (25, 28). ϕ = ΔΔG^≠^/ ΔΔG^ground^, where ΔΔG^≠^ = change in the transition state barrier heights and ΔΔG^ground^ = change in the ground state free energy values. ϕ value indicates the conformational resemblance of a local structural element with either the C or O states at the global transition state (29). ϕ-value analysis has been used as a kinetic ruler to measure the progress of allosteric conformational changes happening in sequence from the TBS to the gate (30, 31). Accordingly, a structural dynamic map showing conformational wave propagation from the TBS to the gate has been proposed in the AChRs in the presence of agonists (29).

In GPCRs (32), nuclear receptors (33) and enzymes (34), allosteric communication has been studied by identifying key side chain residues using molecular dynamic simulations (MD) or from experimental methods including NMR (35), DEER (36), and FRET (37). In AChRs, a principal pathway in the allosteric communication network has been proposed based on electrophysiology experiments (38). Further, salt bridges (39), hydrogen-bond network (40) and structural water network (41) have been surmised to be important in allosteric communication. Further, by MD simulations, it was shown that forced closure of the C-loop led to distinct conformational changes in the receptor and finally, opening of the distant pore of the ion channel in AChRs (42). However, the role of an amino acid residue in receptor activation can be best investigated by strategic mutagenesis and free energy measurements. In neuromuscular AChRs, the free energy changes and ϕ values have been measured for >100 residues across the receptor (30).

Here, by using a combination of single channel patch-clamp experiments, free energy calculations, ϕ-value analysis and structural dynamics simulations, we try to provide insight into the allosteric communication in AChRs. We present the phi map of the unliganded AChR and compare the ϕ and range energy values (see below) with the di-liganded AChRs. We have used the information about ϕ and free energy changes in the liganded vs unliganded receptors to understand the activation barrier at the transition state. Further, we have studied the structural dynamics of the unliganded AChRs to understand the initiation and communication of allosteric signals in the receptor. Our results suggest the following: (1) C-loop capping at the binding site triggers allosteric signal independent of the presence or absence of an agonist at the binding pocket, (2) Activation of the unliganded AChRs proceeds by 2 parallel gating pathways: a predominant pathway that corresponds to the free energy change between C and O (ΔG_0_^mut^) in the presence of a mutation, b. A minor gating pathway, independent of the ΔG_0_^mut^ value of the mutation, that is initiated by the capping of the C-loop at the TBS. We propose that the minor gating pathway is a pre-existing allosteric communication pathway, which is preferentially chosen by the receptor in the presence of an agonist at the binding pocket. (3) Allosteric communication in the unliganded AChRs was mediated by a few high-energy ‘hub’ residues whereas in the liganded receptors, a ‘live wire’ like energy cascades down a sequence of amino acids from the TBS to the gate in the pore.

## Materials and methods

### Cell culture and mutagenesis

Human embryonic kidney (HEK293T) cells were cultured in Dulbecco’s minimal essential medium (DMEM), supplemented with 10% fetal bovine serum and 1% penicillin-streptomycin, at 37°C and 5% CO_2_. AChRs were expressed in HEK cells by transiently transfecting cells with a total of 2μg subunit cDNAs in a ratio of (α1)_2_:b:d:e per 35-mm culture dish using calcium phosphate precipitation method (43). Mutations were introduced using the QuikChange site-directed mutagenesis kit (Agilent technologies), and were confirmed by nucleotide sequencing. Most of the electrophysiological experiments were conducted ∼24 hours after transfection.

### Patch-clamp electrophysiology

Single-channel currents were recorded at room temperature (∼23°C) using the cell-attached patch configuration. The cells were bathed in a K^+^ ringer solution containing (in mM) 142 KCl, 5.4 NaCl, 1.8 Cacl_2_, 1.7 MgCl_2_ and 10 HEPES/KOH, pH 7.4. Patch pipettes were filled with Dulbecco’s phosphate-buffered saline (PBS) comprising (in mM) 137 NaCl, 0.9 CaCl2, 2.7 KCl, 1.5 K_2_HPO_4_, 0.5 MgCl_2_ and 8.1 Na_2_HPO_4_, pH 7.3 adjusted with NaOH. In liganded experiments, agonists were added to the pipette solution from a stock solution to achieve desired concentrations. Patch pipettes, made from borosilicate glass, were coated with sylgard (Dow Corning) and fire-polished to a resistance of approximately 10 MΩ when filled with pipette solution. Single-channel currents were recorded using dPatch low-noise digital amplifier (Sutter Instruments, Novato, CA) at 10 kHz and digitized at a sampling frequency of 50 kHz. Pipette holders and pipettes used in experiments without ligands were never exposed to agonists.

### Estimation of rate constants

Kinetic analyses were conducted by directly fitting kinetic models to idealised single-channel current data using QUB software suite (44). To estimate rate constants, clusters of single channel activity, bordered by non-conducting periods of at least 20 ms, were manually selected. These clusters were digitally filtered at 10 kHz and then idealized into intervals using a segmental K-means (SKM) algorithm (45). A simple kinetic scheme with a C and O (C↔O) state was used to directly fit the idealized event list using a maximum interval likelihood (MIL) algorithm with an imposed dead time of 50 μs (46). We progressively improved our kinetic model by sequentially adding extra C and O states to the basic two-state C↔O scheme, until the maximum interval likelihood (MIL) score improved by <10 units (as per AIC/BIC). The use of a 10-unit criterion was specifically adopted by us to introduce rigor to our single channel data analysis. Ratio of the forward (*f_n_*) and backward (*b_n_*) rate constants associated with the predominant states (see below) was used to calculate the gating equilibrium constant (E_n_), E_n_=*f_n_/b_n_*. The free energy change associated with gating isomerization (kcal/mol) is ΔG_n_ = - 0.59×ln*(E_n_). We could reliably estimate rate constants from intracluster intervals ranging from approximately 10 to 10,000 s^-1^.

#### Estimation of unliganded gating kinetics-choosing a kinetic model

The kinetics of unliganded gating activity is complex (47), with multiple exponential components in the dwell-time distributions. Different models with progressively increasing complexity were fit to the data before choosing the most appropriate model for rate determination. Models were discriminated based on the LL scores and their ability to describe kinetic mechanism. A four-state O_G_↔C_1_↔C_2_↔O_L_ kinetic scheme (parallel model; see Table S1) best described the complexity of unliganded gating (O_G_ and C_1_= predominant gating components (∼90%); O_L_ and C_2_= minor components in the open and shut dwell-time distributions) in most cases though in some cases an additional O state was needed. The ratio of the *f_0_* and *b_0_* associated with the C_1_↔ O_G_ arm was used to calculate the unliganded gating equilibrium constant (*E_0_*). This was used to calculate E_0_ and ΔG_0_ referred to as the major gating (*C_1_↔O_G_*). The opening and closing rates (*f_0_’* and *b_0_’*) determined from the minor long component, were used to calculate E_0_^’^ and ΔG_0_^’^ referred to as the minor gating (*O_2_↔C_L_*). We regularly cross-validated our rate constant estimates from model fits by additionally estimating *f_0_* and *b_0_* from the inverse of the time constants (τ) associated with the shut and open components, respectively.

To further investigate the origin of constitutive activation due to either a parallel vs sequential transition state pathway, we simulated single channel currents using an uncoupled parallel or linear kinetic models (see Table S1, Simulation data not shown). Qualitatively, the data suggest that the simulated single channel data generated by using a linear model do not resemble the constitutive activity seen in the case of AChRs, whereas the simulated data from a parallel model had the characteristic burst behaviour and dwell-time distributions seen in constitutive gating.

To further understand the heterogeneity in the gating of the unliganded receptor, a comparative kinetic analysis was performed using data obtained from liganded experiments involving cholinergic agonists such as choline, TMA and ACh and non-cholinergic agonists such as betaine, NH_4_Cl, nicotinamide and piperidine carboxylic acid. It may be noted that in the latter experiments, even if very high concentrations of the non-cholinergic agonists were used, we cannot be sure if the TBS is fully saturated by the respective agonists.

#### Protein Engineering

Wild-type (WT) AChRs rarely open without agonists. Therefore, unliganded gating of WT receptors cannot be studied directly. We engineered AChRs harboring gain-of-function mutations to increase the probability of constitutive channel activation (48). We incorporated mutations away from the binding pocket and widely separated from each other to ensure minimal interference in agonist binding and reduce any potential inter-residue coupling (48). In neuromuscular AChRs, residues separated by >16 Å behave functionally independent of each other (49).

To understand the complexity of constitutive channel gating kinetics, we utilized several gain-of-function background mutations with increasing gain-of-function. For example, mutations at αA96 amino acid to ∼10^-1^-10^5^-fold gain-of-function (50) resulting in E_0_ ranging from 10^-7^-10^-2^. To this, we added different combinations of background gain-of-function mutations such as bT456I, dI43Q, eE181T+eL269F, eE181W+eL269F, bL262Q+dL265Q and bV266A+dV269A, thus creating large 2-dimensional matrix where each element is a novel background combination (see SI Table 3). The ΔG_0_ for the mutations used ranges over 9 kcal/mol (SI Table 3).

#### ϕ, range energy (ΔG_range_) and transition state barrier height (ΔΔG^ǂ^)

To understand the transition state energy (TSE) landscape of the liganded v/s unliganded receptor, we estimated 3 quantities from the single channel kinetic rates and gating free energy changes: ϕ, range energy and activation energy barrier at the TS (<u>ΔΔG^ǂ^</u>). ϕ is the slope of a linear free energy relationship (LFER) that intricately links the activation energy barrier and the change in the ground state free energy of chemical reactions in the presence of a perturbant such as a mutation (27, 51). It is estimated from the slope of a a log–log plot of the forward (*f_0_*), C→O rate constant versus the C↔O equilibrium constant (E_0_) (rate–equilibrium free energy (R–E) relationship or REFER) for a series of mutations at any residue. Range energy (ΔG_range_) is theoretically the total possible free energy change (kcal/mol) when an amino acid is substituted for all other 19 amino acids, which provide insight into the local conformational changes during C↔O gating. However, mutating each amino acid with 19 residues is practically not possible. Therefore, to calculate the range energy of a residue at a particular position in the receptor, the residue was typically, mutated to at least 4 to 10 side chains. The effect of the substitutions was calculated as a change in the gating free energy, *ΔΔG_0_*. Range energy for a residue was therefore calculated as:

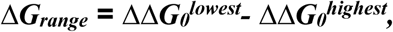

where ΔΔG_0_^lowest^ and ΔΔG_0_^highest^ are the lowest and highest free energy changes associated with a mutation of a particular residue. For the liganded receptor, the values of ϕ and range energy for several residues were collated from the published work of Purohit et al, 2013 (30) and others (24, 39) (SI Table 10).

The change in the transition state barrier height (***ΔΔG^ǂ^***) was calculated for each mutated position, both in the unliganded and the liganded receptor using ϕ and range energy (***ΔG_range_***). Assuming that the transition state is intermediate between C and O in geometry and topology, ΔG^‡^ can be expressed as a combination of ΔG_C_ and ΔG_O_ as:

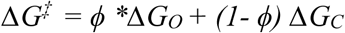

where 0≤ ϕ ≤ 1. For any perturbation, equation 2 can be re-written as:

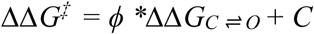

Where C is a constant whose value is ΔG^‡^_0_, which is the intrinsic activation energy barrier and ΔΔG_C ⇌ O_ is the ground state energy which is the range energy for a particular residue. So, the barrier height was calculated as:

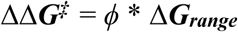

The values obtained for *ϕ, ΔΔG_0_* and *ΔΔG^‡^* were substituted in the B-factor column of the PDB ID: 9AWJ to generate structure maps of *ϕ,* range energy and transition state barrier height for the unliganded and the liganded receptor.

#### In silico mutagenesis and MD simulation

Recently solved ACh bound structure of *Bos taurus* (PDB Id: 9AWJ) was used for the present study. We strategically incorporated the following mutation: αG147A, αP197A, αG153K, and eE181W using the Structure Editing module of UCSF Chimera v1.16 (52) in the receptor. ACh was stripped from both the binding pockets of all the systems. A total of five systems were prepared: WT receptor and the four point mutations. The Dunbrack 2010 rotamer library (53) was used to select the orientation of the mutated residues with the least atomic clashes. The energy of the systems was minimized by 1000 steps of steepest descent with the step size of 0.02 Å.

The individual WT and mutated structures were embedded in a bilayer membrane that consists of 1-palmitoyl-2-oleoyl-sn-glycero-3-phosphocholine (POPC) of size 150Åx150Å using the Membrane builder plugin of VMD v1.9.4 (54). The membrane-embedded systems were solvated with TIP3P water (55) and ionized using 150 mM NaCl. The lipids and water in the vicinity of the membrane-embedded protein (within 1.5Å) were removed. The MD simulation was performed using NAMD v2.14 (56) with CHARMM36m as the force field (57).

Initially, all prepared systems were subjected to 50000 steps of energy minimization via the steepest descent method, followed by an equilibration of 20 ns by gradually reducing the harmonic restraints applied on the lipid molecules. Subsequently, all systems were simulated without any restraints for about 300 ns (varying with systems) under NPT conditions using standard periodic boundary conditions and minimum image convention. Periodic conditions were applied, and the Langevin dynamics method (58) was used to set the temperature at 310 K and pressure at 1 atm. Using the SHAKE algorithm (59) an integration timestep of 2 fs was used while keeping all the covalent bonds involving hydrogen atoms constrained to their equilibrium bond length. The particle mesh Ewald algorithm (60) was used to calculate electrostatic interactions. A switching distance of 10Å and a cutoff distance of 12Å were applied for non-bonded interactions. The simulations were run at a timestep of 2 fs, and the coordinates were saved every 100 ps for subsequent analyses. All other parameters were left to the default values. The stability of the MD simulations was confirmed by plotting the root mean square deviation (RMSD) of the protein over the course of the entire trajectory.

#### Structural analysis

PyMol v2.5.7 was used to visualize and extract images (https://pymol.org). The system stability was analysed using RMSD plots. All structural analyses were done from the last 100 ns of the production run using VMD v1.9.4. SigmaPlot 11.0 was used for plotting the data and curve fitting (*Sigma Plot, Version 11. San Jose, CA*).

To correlate the structural dynamics of the loop C capping in simulations with the experimental single channel data, the position of the loop C was determined by quantifying the distance between the residues aC192 (loop C) and aW149 (loop B). The average distances between aC192 and aW149 were estimated by plotting distribution histograms and fitting it with a gaussian function. The corresponding residence time of each conformation was calculated over the trajectory using Markov State modelling (61). The MSM analysis utilized the Gaussian Mixture Models (GMM) for metastable state identification from protein conformational dynamics. The algorithm performed optimal state determination via AIC/BIC and clustering validation (silhouette scores), testing n-state models from 2 to 8 states. State assignment utilized GMM soft clustering on distance trajectories, computing probability densities P(x|θᵢ) for each state i with parameters θᵢ. The core MSM construction calculated transition probability matrices T(τ) at lag time τ by counting state-to-state transitions:

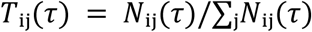

where Nᵢⱼ(τ) represented transitions from state i to j after time τ. Optimal lag time selection employed timescale convergence analysis, identifying *τ* where implied timescales stabilized, satisfying the Markovian assumption. Chapman-Kolmogorov validation tested MSM quality by comparing predicted v/s observed transition probabilities at different time intervals, ensuring accurate kinetic modelling of loop C conformational transitions.

The experimental data obtained from WT and mutants (aG147A, aG153K, aP197A and εE181W) were used to calculate the state residence probabilities of the major and minor gating components. The estimation of state residence probability was done using the formula:

> *τ (mean time constant) * Area under a curve from the open dwell time distribution.*

To compare the data obtained from simulations and experiments, a correlation scatter was plotted between the residence probability obtained from experiments and the ones obtained from simulations. The distribution was fitted using a linear regression model and the R^2^ value was calculated to determine the goodness of the fit.

## Results

### Unliganded gating of neuromuscular AChRs

#### Comparison between liganded vs unliganded gating kinetics

Figure 1b shows the representative single channel current traces and the dwell-time duration distribution histograms for the WT AChRs in the presence of saturated concentrations of Cho (20 mM), TMA (1 mM) and ACh (500 μM). The single channel kinetics were simple, described by single exponential probability density functions. AChRs in the absence of neurotransmitter rarely opens. To compare the single channel activities of liganded vs unliganded AChRs, we engineered gain-of-function mutations that resulted in constitutively active receptors with *Po* values comparable to that of the liganded AChRs. For example, a mutation at αA96H in loop A in the extracellular domain shifts the equilibrium constant (E_0_) by ∼100,000 fold (50). Addition of another mutation βT456I in M4 further increases the E_0_ by ∼2-fold. Therefore, the expected E_0_ in the presence of the combination of αA96H+βT456I ≈ 0.15 (7.4×10^-7^×100000×2 = 0.15; see Methods). Fig. 1c shows the representative single channel currents and dwell-time duration distributions from AChRs having (αA96H+βT456I) mutations in combination. The observed E_0_ for this background =0.064. This is comparable to the di-liganded gating equilibrium constant (E_2_) for choline, a partial agonist of AChRs (Fig. 1b). Then, we introduced background gain-of-function mutations to design constitutively active AChRs which matched in their E_0_ values to the E_2_ of TMA (E_2_^TMA^=2.4 vs E_0_ for [αA96N + βT456I + εE181T + εL269F] = 1.6; see Methods) and ACh (E_2_^ACh^=25.4 vs E_0_ for [αA96L + βL260Q + δL265Q] = 20.5). Though the gating equilibrium constants for the liganded and unliganded AChRs were comparable, the kinetics were different and more complex and heterogeneous in the unliganded AChRs (Fig. 1c vs 1b). The complex gating of unliganded AChRs have been previously documented for several background mutation combinations by us and others (31, 47).

Surprisingly however, the binding of unconventional non-cholinergic agonists to AChRs displayed complex kinetics, analogous to the unliganded AChRs. Fig. 1d shows the representative current records and histograms from WT AChRs in the presence of Glycine-Betaine (10 mM), NH_4_Cl (100 mM), Nicotinamide (5 mM) and Piperidine carboxylic acid (1 mM). The E_2_ values for these agonists range from 2.2×10^-5^-0.004.

#### Allosteric signal ‘initiation’: Insights from unliganded gating complexity

Fig. 2a shows the representative single channel currents, dwell time duration distribution histograms and apparent E_0_ values obtained from AChRs having only 1 mutation. At E_0_ < 0.002, the channel open time distribution histograms could be described by a single exponential component. With increasing gain-of-function, we observed the evolution of a ‘minor’ component in the open time distribution histograms. In αA96H, the kinetics of the clustered single channel data could be best described by a kinetic model comprising of 2 closed (C) and 2 open (O) (SI Table 3; see Methods) states. Then we incorporated combinations of gain-of-function mutations in the receptors to understand the role of underlying structural element(s) in the unliganded gating heterogeneity. In general, for mutations away from the binding site, (at the hydrophobic gate in the pore for e.g.) the kinetics of unliganded gating could be explained by 2C and 2O states (O_G_-C_1_-C_2_-O_L_, see below; Fig. 2b, right; SI Table 3; see Methods). Of the 2 open states, 1 was a major component (90±10%) and the other was a minor component (10±3.4%). The minor component manifested with relatively longer lifetime (τ) for background mutation combinations with E_0_ values ranging between ∼0.01<E_0_<20. However, for E_0_ > 20, the long-open component was not observed (Fig. 2b, bottom), presumably superimposed by the predominant gating component. Fig. 2c shows single channel currents, dwell-time histograms for mutation combinations having E_0_ > 20 (ΔG_0_<-2.0 kcal/mol). For different combinations with increasing E_0_ (SI Table 3), we observed a simple C↔O kinetics, akin to the gating observed in the presence of cholinergic agonists (For reference, ΔG_2_^ACh^ = - 1.9 kcal/mol). We will refer to the major component as the gating component (O_G_) and the minor one as the long open component (O_L_), hereafter.

**Figure 2.**
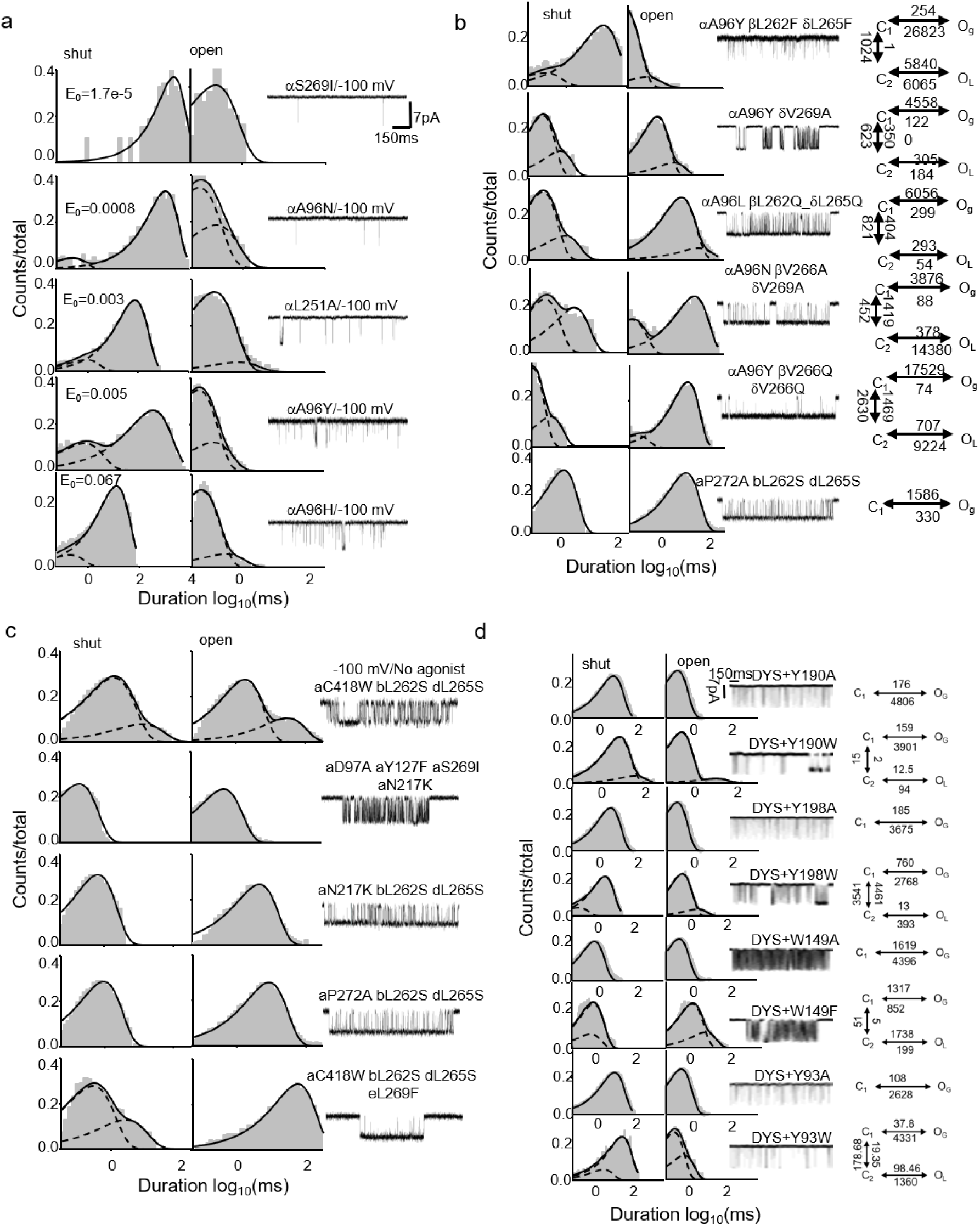
Evolution of minor gating component in unliganded gating of AChRs. **(a)** Origin of kinetic complexity in AChRs containing a single amino-acid substitution (with increasing E_0_). Representative single-channel current recordings and dwell-time distributions are shown for unliganded AChRs containing individual amino-acid substitutions at the positions indicated in the figure. Mutant receptors with an intrinsic gating equilibrium constant, (E_0_)< 0.02 exhibit relatively simple C–O kinetics. Note the progressive evolution of an additional minor exponential component in AChRs with (E_0_ > 0.02) in the open-duration distribution. **(b)** Kinetic complexity of unliganded AChRs containing mutations in different structural elements. Representative single-channel current records, dwell-time distributions and kinetic models for AChRs containing combinations of background mutations located at different positions throughout the receptor. The dwell-time distributions consistently show two shut- and two open-time exponential components. In all cases, the kinetic complexity is best described by the (O_L_–C_2_–C_1_–O_G_) kinetic model (see SI Table 1). **(c)** Simple C–O kinetics in unliganded receptors with E_0_ > 30. Representative single-channel current records and dwell-time distributions for unliganded AChRs containing combinations of gain-of-function background mutations that produce receptors with *P_o_*>0.99. In each case, the activity is adequately described by simple C–O kinetics. **(d)** Relationship between unliganded kinetic complexity and the agonist-binding site. Representative single-channel current records and dwell-time distributions shown for AChRs containing mutations at the agonist-binding site including W149, Y190, Y198, and Y93. Substitution of the aromatic residue with Ala eliminated the minor gating component. In contrast, aromatic-to-aromatic substitutions retained the long open-time component observed in the corresponding background receptors.

##### Minor gating and the binding site

Purohit and Auerbach ((47)) have shown that mutations of the aromatic residues at the core binding pocket (αW149, αY198, αY190, αY93) eliminated long openings in the unliganded gating. However, we observed that though most of the mutations indeed eliminated the minor long openings, there were clear exceptions. Aromatic → aromatic substitution including aY190W, aY198W, aW149F, and aY93W did not eliminate the O_L_ component (Fig. 2d). This suggests that a TBS constituted by 5 aromatic residues (even in the mutated AChRs) is critical to the origin of the minor gating component. A non-aromatic substitution in the TBS presumably alters the binding site conformation that does not support the minor gating.

In loop B of the α-subunit, αG147 residue has been proposed as an “activation hinge” that determines the flexibility of the loop and is associated with high ΔΔG_0_ values (Purohit and Auerbach, 2011). On the other hand, αG153 is proposed as a “deactivation hinge” that has opposing action. We investigated the kinetics of these two residues on a gain-of-function background (αD97A+αY127F+αS269I; SI Table 3). Fig. 3a shows representative single channel currents, dwell-time duration distribution histograms and kinetic model for the background αD97A+αY127F+αS269I. The dwell-time durations were well-described by the O_G_-C_1_-C_2_-O_L_ scheme, as above. But, in the presence of αG147A mutation on this background simplified the unliganded gating behaviour and the kinetics could be described by a simple C_1_↔O_G_ scheme. In contrast, αG153A/K mutations on the same background exaggerated the complexity of the unliganded gating, which could only be described by adding 1-2 extra O state(s) (Fig. 2d). Interestingly, in the presence of both the (αG147A+ αG153A) mutations, we observed a simplified C_1_↔O_G_ kinetics. This indicates that mutations at the glycine hinge residues, which are known to alter the agonist binding affinities (62), reciprocally regulate the heterogeneity in the unliganded gating.

**Figure 3.**
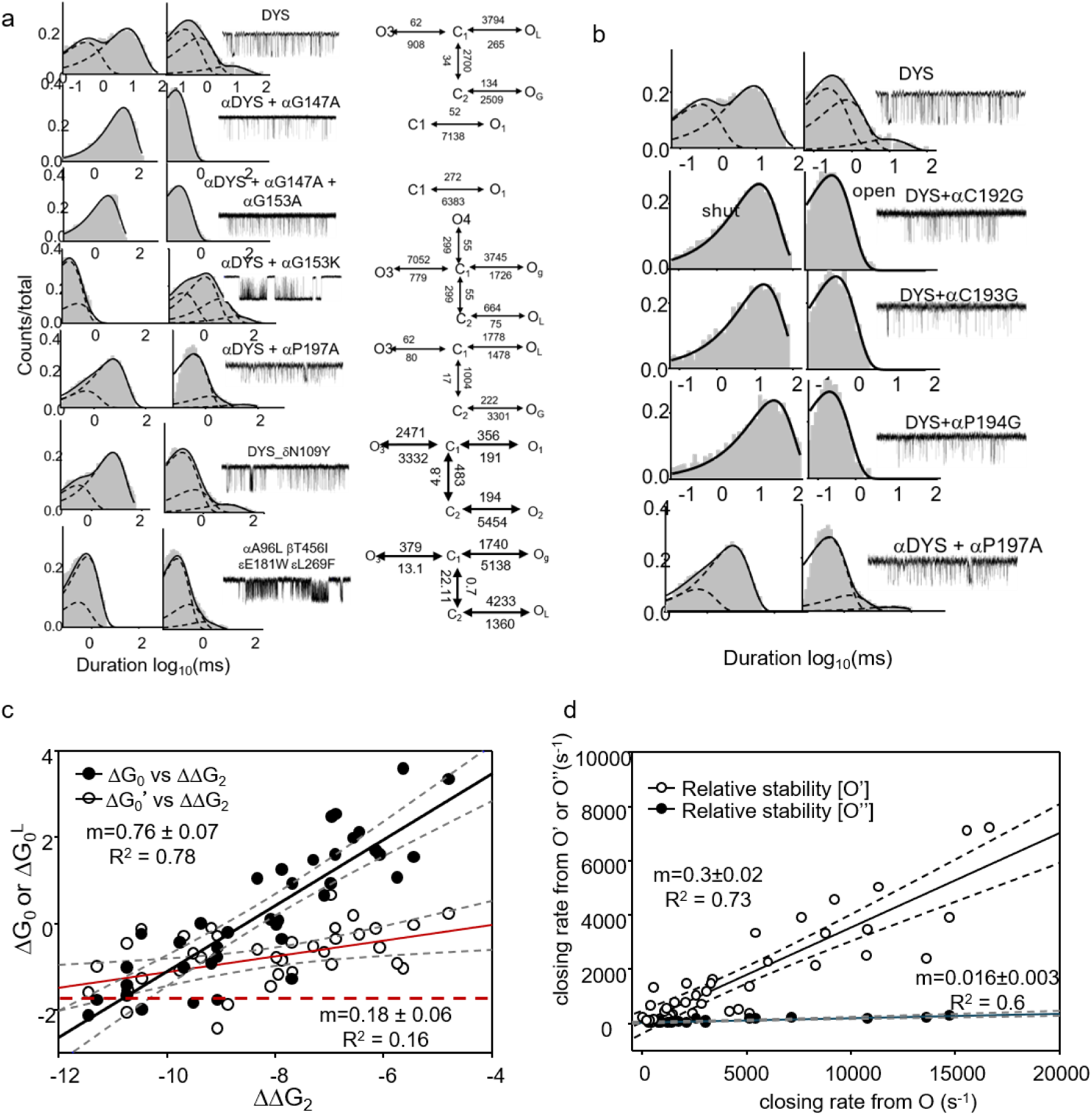
Parallel gating pathways in unliganded AChR activation. **(a)** Relationship between the residues near the agonist-binding site and gating pathways of unliganded gating. Representative single-channel current recordings, dwell-time distributions and kinetic models shown for the glycine-hinge residues αG147 and αG153 in the presence of gain-of-function background mutations (shown in Figure), and for mutations at aP197, δN109, and εE181. Note the differential modulation of the minor gating component by mutations in the vicinity of the binding site. **(b)** Effects of loop C mutations on the minor gating pathway. Representative single-channel current records and dwell-time distributions obtained from AChRs containing glycine substitutions at residues C192–P197 of loop C. The substitutions exert contrasting effects on the minor gating component, indicating that individual loop C residues differentially influence the parallel unliganded gating pathway. **(c)** Correlation between expected (ΔΔG_2_^mut^) and experimentally estimated free-energy changes associated with the major (ΔΔG_0_) and minor (ΔΔG_0_^’^) gating components. Scatter plots show the relationship between the expected and experimentally estimated mutation-induced free-energy changes. Solid lines represent linear regressions through the data. The corresponding goodness-of-fit parameters are shown in the figure. Dotted red-line = ΔG_2_^ACh^**. (d)** Correlation between the relative stabilities of the major and minor gating components. Scatter plots show the relationship between the stabilities of the major and minor gating components. Solid lines represent linear regressions through the data, and the corresponding statistical and goodness-of-fit parameters are shown in the figure.

Loop C at the TBS is a key determinant of agonist binding (63–65). We studied the unliganded gating for some of the loop C residues including αC192, αC193, and αP194. In the presence of Gly mutations at these positions resulted in a simplified C_1_↔O_G_ kinetics like the loop-B αG147 residue mutations (Fig. 3b). On the flip side, mutation at the neighbouring αP197 (Ala) led to heterogeneous channel kinetics with >2 O states. This indicated that mutations at residues in the loops B and C, within 8Å of the core binding pocket, alter the channel kinetics analogous to the core aromatic residues at the binding pocket. Further, we investigated the kinetics of AChRs in the presence of mutations at residue εE181 and δN109 (63, 66, 67) on the complimentary side of the TBS which have been implicated in agonist binding. Mutations at these sites enhanced the heterogeneity of constitutive gating and resulted in up to 3 O states (Fig. 3a, lower)., exaggerated the occurrence of the minor long component or led to appearance of additional longer O components or both (Fig. 3). Their kinetics could not be described by the O_1_-C_1_ ─ C_2_-O_2_ model but required an additional O state. The unliganded gating kinetics, the *f_0_*, *b_0_*, *f_0_’*, *b_0_’*, E_0_ and E_0_’ values for all the engineered AChRs are given in SI Table 3. The above results suggest that mutations of amino acid residues, implicated in regulating agonist binding, at and near the TBS alter the minor gating component.

##### Parallel gating pathways in the unliganded AChRs

There are contrasting reports on whether AChRs, in the presence or absence of agonists, follow the same (47, 68) or multiple activation pathways (69). We hypothesized that if the activation of AChRs proceeds along 1 gating transition pathway in the TSE, then, energetic perturbations such as mutations should result in equivalent or correlated changes in the O_G_ vs O_L_ gating components. Fig. 3c is a scatter plot of the expected ΔΔG_2_^mut^ (-0.59*ln[E_2_^mut^/E_2_^WT^]) in the presence of combinations of gain-of-function mutations (see Methods) vs the observed ΔG_0_^obs^ estimated from the predominant gating component or the minor gating component (ΔG_0_^L^). The scatter plots were fitted with linear regressions. The slopes of the linear regressions were 0.76±0.07 (R^2^=0.78) and 0.18±0.06 (R^2^=0.17), respectively. This indicates that the ΔG_0_^obs^ values associated with the predominant gating component correlates with the sum of energetic contributions from independent mutations whereas the ΔG_0_^L^ values are mostly independent of the background mutations. Earlier, it has been shown that the predominant gating component follows predictably the thermodynamic cycle (within the cycle;(8)) and there is a linear correlation between the ΔΔG_0_ and ΔΔG_2_ (30, 70, 71) values with a slope of 1. The average ΔG_0_^L^ values (-1.14±0.65 kcal/mol; n = 38) approached the diliganded gating energy, ΔG_2_ = - 1.9 kcal/mol (red dotted line in Fig. 3c).

In some cases, >2 O states were needed to describe the kinetics of unliganded gating (Fig. 3a). To compare their kinetics with the frequently observed [O_G_-C_1_-C_2_-O_L_] type of kinetics, we plotted a relative stability plot (Fig. 3d), which describes the correlations between the closing rates from O_G_ (*b*) vs O_L_ (*b’*) and O_G_ (*b*) vs O_L_’’ (*b **’***). The data were fitted by linear regression lines (0.3±0.02 (R^2^=0.73) and 0.016±0.003 (R^2^=0.6)). These results indicate that the minor gating components of unliganded receptors neither kinetically (different rates) nor energetically (different ΔΔG0’) resemble the predominant gating pathway of the receptor.

##### Loop C capping and initiation of allosteric communication

To understand the effect of the above binding site mutations in minor gating, we studied the structural dynamics at the TBS by performing all atom molecular dynamics simulation (MD) using the wild-type and mutant AChRs (PDB ID 9AWJ). Capping movement of the loop C (displacement towards the aromatic binding pocket) has been implicated in the low-affinity binding of cholinergic agonists in the AChRs (39, 72) and more recently, shown during low (AC) → high (AO) affinity state transition (73). In total, we prepared 5 different systems (see Methods for details). The systems were unliganded WT, aG147A, aG153A, aP197A and eE181W. The systems were prepared in POPC membrane, saturated with water and ions, ligands were stripped, equilibrated for ∼20 ns and production runs were performed for 120 ns each in triplicates (see Methods). The most conspicuous change between the WT vs the mutant receptors was the position of the loop C. Fig. 4a shows the C-loop position in the average structures obtained from the simulations.

**Figure 4.**
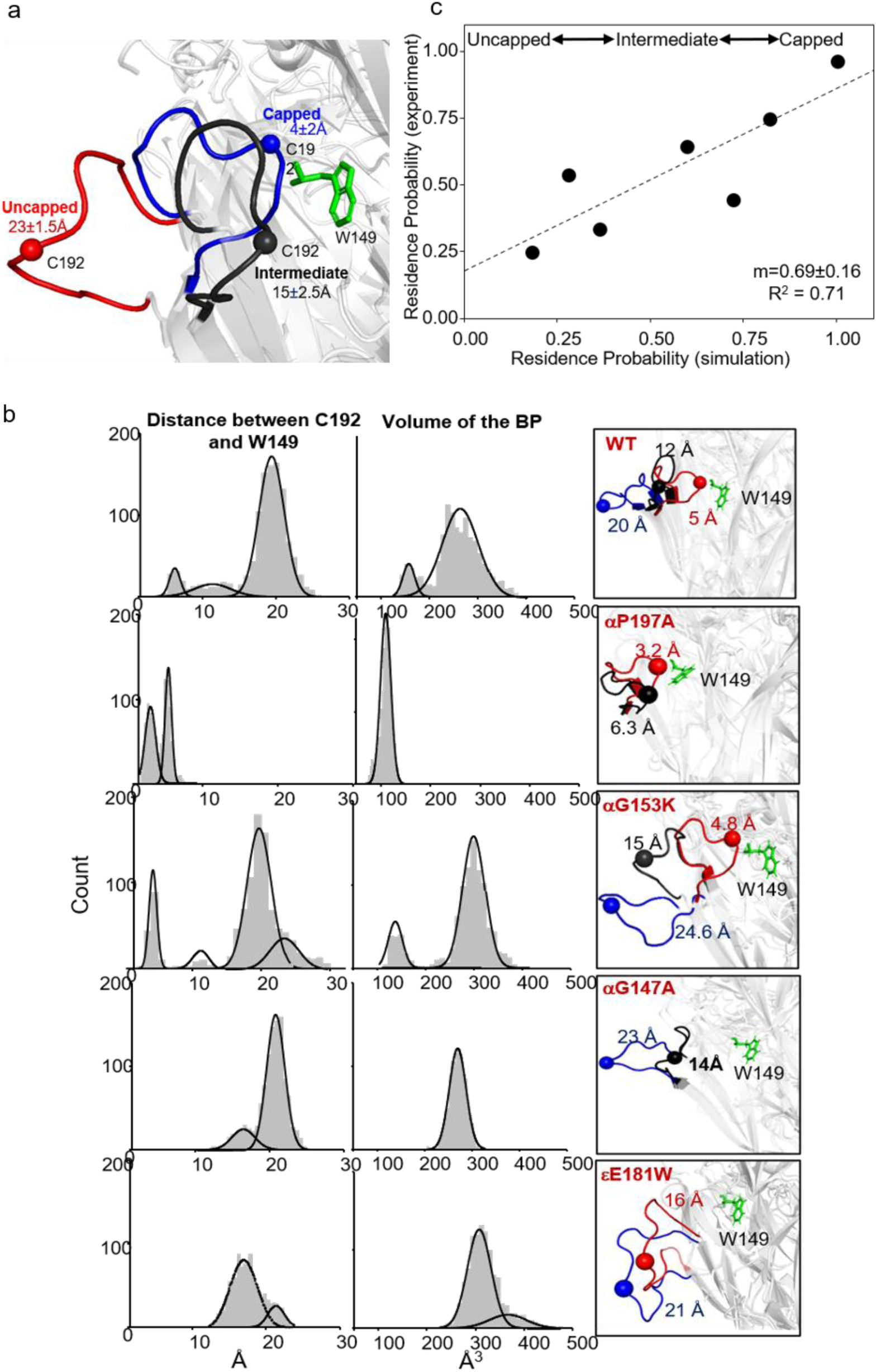
Loop C as a structural trigger for allosteric communication at the agonist-binding site. **(a)** Distinct conformational states of loop C in the unliganded AChR. Zoomed in view of the neurotransmitter-binding pocket (αW149 shown to demark the TBS) in the agonist-stripped AChR structure (PDB: 9AWJ), showing the position of loop C relative to the aromatic binding pocket. Three distinct loop C conformations are evident: 1. conformation positioned closest to the aromatic binding pocket (capped), 2. conformation located farthest from the pocket (uncapped), and an intermediate conformation (intermediate-capped). Average distances of the C-loop for the different conformations are shown in the figure. **(b)** Structural analysis of loop C distance and agonist-binding-pocket volume in WT and mutant AChRs. Left, distributions of the distance between αC192 and αW149 are shown for the indicated mutations. Right, distributions of the agonist-binding-pocket volume are shown for the mutant and wild-type receptors; pocket volumes were calculated as described in the Methods. The distributions associated with the αG153 and αP197 mutations exhibit greater conformational complexity than those observed for the αG147 mutations. **(c)** Correlation between experimental and simulated loop C residence probabilities. Scatter plot showing the relationship between the residence time of the loop C conformational states calculated from Markov state model analysis and the corresponding residence probability estimated from experimental single-channel dwell-time distributions. The solid line represents a linear regression fit to the data. The goodness-of-fit and associated statistical parameters are shown in the figure.

To quantify the C-loop positions we measured the average distance between loop C (aC192) and B (aW149) (Fig. 4b). In the WT, the C-loop occupied 3 distinct conformations which were at 21.9±1.99 Å (∼70 % occupancy, I), 16.6±1.2 Å (10% occupancy, II) and 11±0.8 Å (20% occupancy, III) away from the aW149 residue in the B-loop. We define the conformations I, II, and III as ‘uncapped’, ‘intermediate’ and ‘capped’ respectively. In contrast, in the αG147A mutant, the position of the C-loop was predominantly in the farthest uncapped conformation (∼99% occupancy). However, in the αG153K mutant, the C-loop samples all the 3 conformations though it remains in the capped conformation for substantial amount of the time (∼60%). We then measured the volume of the binding pocket for the WT and αG147A/αG153K mutants. The binding pocket volume distributions for the mutants were similar as above (Fig. 4b, right). This indicates that the contrasting effects of the Gly hinge residues on the minor gating components (observed above) might be correlated to the opposing conformational dynamics observed in MD simulations for these mutations. C-loop occupied 3 distinct conformations in the simulation of αP197A mutant AChRs, which further corroborated the exaggerated minor gating component observed in experiments (Fig. 4b).

To further understand the correlation between C-loop conformational dynamics and the emergence of functional major and minor gating, we correlated the state residence probabilities obtained from the MD simulations of WT and mutant AChRs with the single channel state residence probabilities obtained from mean open time constants and the area under the major and minor components in dwell-time duration distribution histograms. The hypothesis is that a capped C-loop conformation results in long minor gating whereas uncapped/partially capped conformations result in the predominant major gating. The state transition probabilities for uncapped ↔ intermediate ↔ capped conformations from simulations were estimated by using Markov State modelling (61). Fig. 4c shows the correlation scatter plot fitted with linear regression. The slope of the regression line was ∼0.69±0.1 (R^2^ = 0.71). This suggests that the residence probability of the major gating component was correlated with the uncapped or intermediately capped C-loop conformation, whereas the residence probability of the minor component was correlated well with the capped conformation of the C-loop. Since, the ΔG_L_ (free energy of the minor long opening) and ΔG_2_ (liganded gating energy) values are comparable and C-loop capping is obligatory for liganded gating, we propose that in the unliganded AChRs the minor long gating represents a pre-existing pathway which is preferentially chosen in the presence of the agonists in the liganded receptor.

#### Allosteric communication in unliganded vs liganded AChRs: Phi, Range energy (ΣΔG_0_) and ΔΔG_0_^‡^

To understand the position of the transition state along the reaction coordinate, we estimated ϕ-values from the ratio of the change in *f_0_* vs E_0_. For the major and minor gating components, we defined ϕ_1_ and ϕ_2_ as (δ*f_0_/*δ*E_0_*) and *δf_0_’*/*δE_0_’,* respectively. SI Table 8 shows the single channel kinetic parameters, ϕ_1_ and ϕ_2_ values for several amino acids in AChRs in the absence of any agonists. For all the residues ϕ_1_≠ ϕ_2_.

To compare the ϕ values in the unliganded vs liganded AChRs, we collated the ϕ values published Purohit et al, 2013 (30) (see SI Table 9), replotted the distribution and fitted the distribution with gaussian functions (Fig. 5a, left). In the liganded AChRs, the ϕ values showed a multi-modal distribution with at least 5 distinct peaks. They have been construed as the 5 different structural blocks of amino acids which undergo C→O transition at temporally sequential manner (29, 30). The distinction in boundaries was determined statistically by using a k-means algorithm. Fig. 5a, right is the distribution of ϕ values estimated for the unliganded AChRs (Right) in the current work. In the unliganded AChRs, we do not see clear multimodal distribution of ϕ blocks, rather, approximately, 50 % of the residues in the unliganded AChRs have ϕ values in the intermediate range ∼0.6±0.2. Only a few residues at the ECD-TMD interface had a ϕ value close to 1 in contrast with the liganded AChRs, where ∼30% of the residues have ϕ-values >0.8. In the unliganded AChRs, most of the aromatic residues at the TBS have ϕ-values <0.7. To gain structural insight from this analysis, we plotted the distribution of ϕ values as heatmaps in the pdb structure of the apo AChR (9AVV;(63); Fig. 6a). Fig. 6a shows that there are fewer ϕ populations of residues in the unliganded vs the liganded AChR. In the liganded receptor, there was clear and progressive decline of ϕ values from the TBS to the gate, whereas in the unliganded AChRs the ϕ values were relatively more scattered.

**Figure 5.**
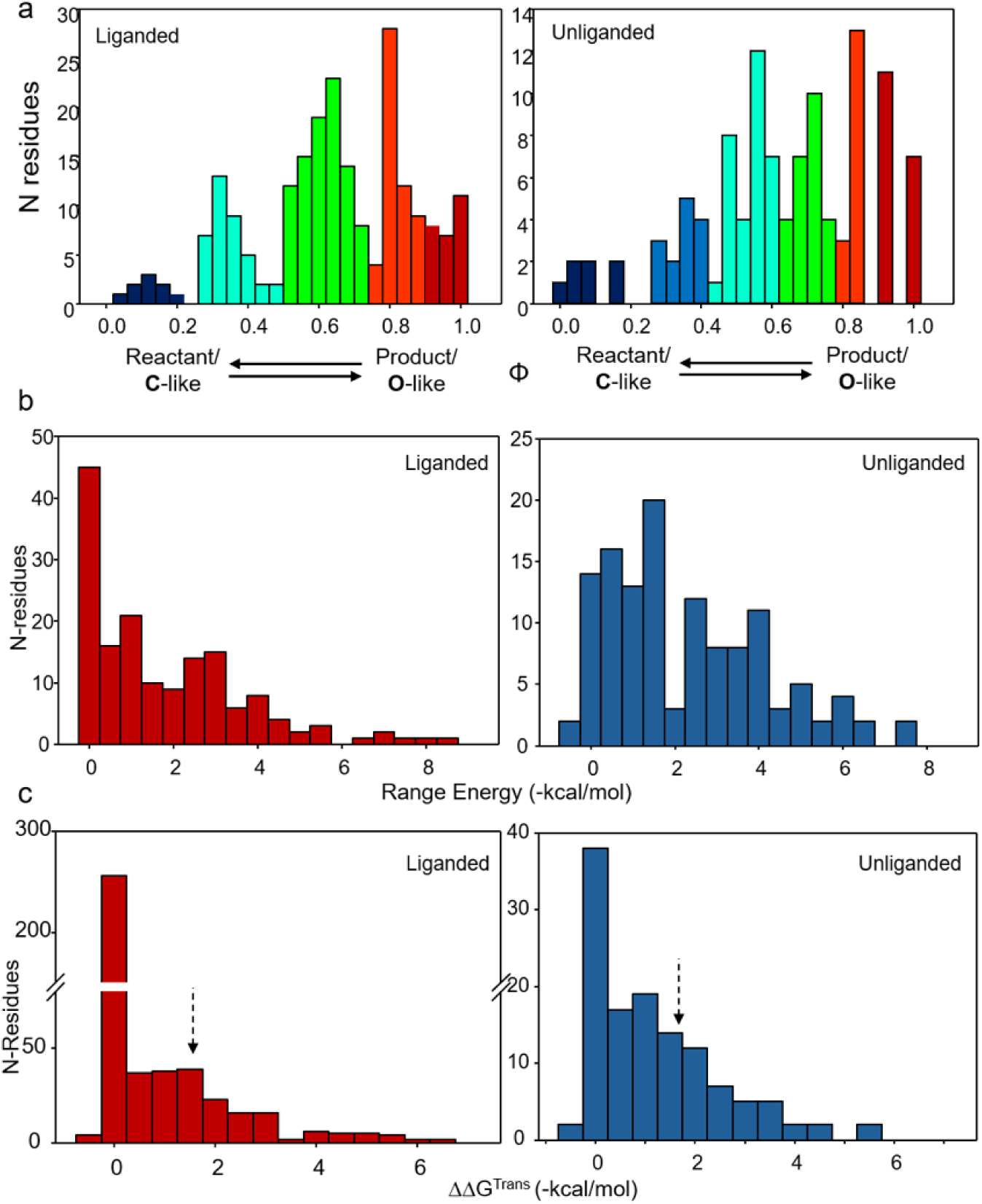
Distributions of ϕ-values, range- and activation energies in unliganded vs liganded AChRs. **(a)** Distribution of ϕ-values in unliganded and liganded AChRs. ϕ-values collated from the literature for liganded AChRs and those estimated in the present study for unliganded AChRs are shown as distribution histograms and colour-coded according to the accompanying heat map. The liganded ϕ-value distribution resolves into approximately four to five distinct Gaussian distributions, whereas the unliganded distribution does not exhibit clearly separated components. **(b)** Range-energy distributions in unliganded and liganded AChRs. Distribution histograms of range energies collated from the literature for the liganded AChRs with those estimated in the present study for unliganded AChRs. Note the marked difference in the overall range-energy distributions of unliganded and liganded receptors. (**c**) Change in activation energy barrier energies (ΔΔG^‡^) for the liganded vs unliganded AChRs (see Methods for calculation) as distribution histograms. Note the stark difference in activation energy changes between the unliganded vs liganded conditions.

**Figure 6.**
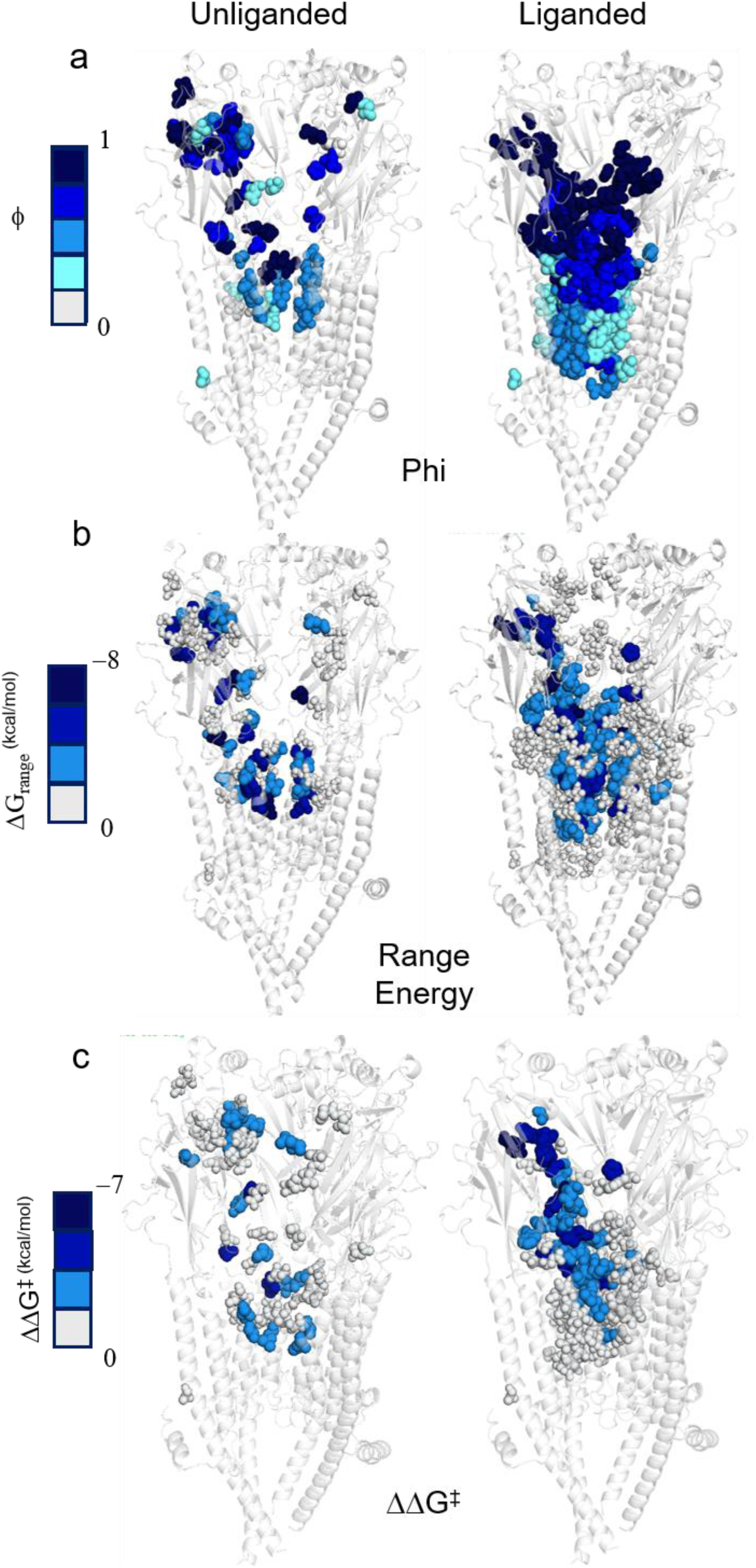
Allosteric communication in liganded vs unliganded AChRs. Structural mapping of (ϕ)-values **(a)**, range energies **(b)**, and activation-barrier heights (ΔΔG^‡^) **(c)** in liganded vs unliganded AChRs. Experimentally estimated or literature-derived ϕ-values, range energies, and ΔΔG^‡^ were mapped onto the AChR structure (PDB: 9AWJ) by replacing the crystallographic temperature-factor values with the corresponding energetic parameters. Residues are colour-coded according to the heat map shown for each structural representation. The unliganded ϕ-value map does not reveal a clear spatial pattern, whereas the liganded receptor exhibits a progressive ϕ-value gradient across the structure. Mapping of ΔΔG^‡^ in the liganded receptor reveals a distinct, approximately linear, live-wire-like allosteric communication network extending from the transmitter-binding site to the channel gate. This pathway is not apparent in the unliganded receptor. Note the presence of a few spatially distributed high-energy hub residues in the unliganded receptor.

To further understand allosteric communication, we compared the energetics of the unliganded vs liganded AChRs. To that end, we measured the ΔΔG_0_ and collated all published ΔΔG_2_ values and estimated the range energy (ΣΔG) for both the unliganded vs liganded AChRs. Side chain substitutions at different positions in the receptor may stabilize, destabilize, or cause no significant change in the O vs C gating equilibrium. Accordingly, mutations cause negative, positive, or insignificant gating free energy changes (ΔΔG_0_) in AChRs. ΣΔG is a minimum approximate estimate of the C↔O gating energy change in their local environment. Fig. 5b (left and right) are the distributions of ΣΔG_2_ vs ΣΔG_0_ respectively, which showed significant differences between them. The values were further mapped onto the pdb structure as above (Fig. 6b), which showed that in the unliganded AChRs, there are only a few sporadically distributed residues which are associated with high-free energy changes. SI Table 8 shows the positions, substitutions, and the range energies for 25 different residues in the ECD and TMD of the AChRs in the absence of any agonists. In our experiments, 2-fold change in equilibrium constant or ±0.4 kcal/mol is considered as statistically significant change in measurement. However, in range energy interpretation, we have used a higher cutoff to identify residue positions showing significant energy changes.

Then, we measured the change in transition state barrier height (ΔΔG^‡^) for each of the positions in the unliganded vs liganded AChRs. ΔΔG^‡^ = ϕ × ΔΔG_0_. SI Table 8 and 9 show the ΔΔG^‡^ values for all these residues. Fig. 5c shows the distribution of ΔΔG^‡^ for the unliganded vs liganded AChRs. To visualize the changes in ΔΔG^‡^ in the receptor during C↔O gating transition, we superimposed the free energy changes of the TS in the pdb structure as above (Fig. 6c). The ΔΔG^‡^ values are represented in the structure by a heat map with cooler colours representing the relatively larger free energy changes. In the unliganded AChRs, some of the residues including, αA96H, αY127F, αP265, and βV266 sporadically appeared in the structure as high-energy ‘hub’ positions, where the activation barrier was greatly lowered by <-3.5 kcal/mol during C↔O transition. In other words, these are the residue positions in an unliganded WT receptor play key role in allosteric C↔O gating conformational change. The other positions that we investigated presented no clear pattern in energy distribution.

In contrast, in the liganded AChR, the distribution of amino acids associated with ΔΔG^‡^ values <-3.5 kcal/mol were highly organized and presented a clear pattern. To visualize the pattern, the residues associated with small energy changes (<-3.5 kcal/mol) were colour muted. In the liganded receptor, the high-energy hub residues (>-3.5kcal/mol) were observed to be form a ‘live wire’ like connection from the TBS to the gate as a (Fig. 6c, Table 9). This suggests that all the residues on this network lower the activation energy barrier to the same level.

##### Allosteric communication: amino acid side chain vs Cα backbone?

To understand whether allosteric communication happens through the side chains or through the C-alpha backbone of the protein, we analysed the ΔΔG values for Gly/Pro→X (any mutation) and X→ Gly/Pro mutations from our current and other previously published work (see Methods). SI Table 6 and 7 show all the free energy changes for the Gly/Pro→X and X→ Gly/Pro mutations. Fig. 7a and b are the distributions of ΔΔG_0_ values for these substitutions. The energy distributions are not significantly different from each other within ±2 kcal/mol range (shown by dotted lines). In the X→ Gly/Pro mutations or in other words, the energy changes associated with Gly/Pro substitutions at any position could be explained by simple local resettling energy during unliganded C↔O gating. However, for a few strategically placed Gly/Pro residues, including αG147, αG153 at the binding site and αP265, αP272 at the ECD-TMD interface (SI Table 6 & 7), substitutions to other residues were associated with significantly high ΔΔG_0_. This indicated that perhaps, in the unstructured loop regions in the receptor C-alpha backbone might significantly contribute to allosteric signalling, whereas in the rest of the receptor, side chain resettling predominantly drives allosteric communication.

**Figure 7.**
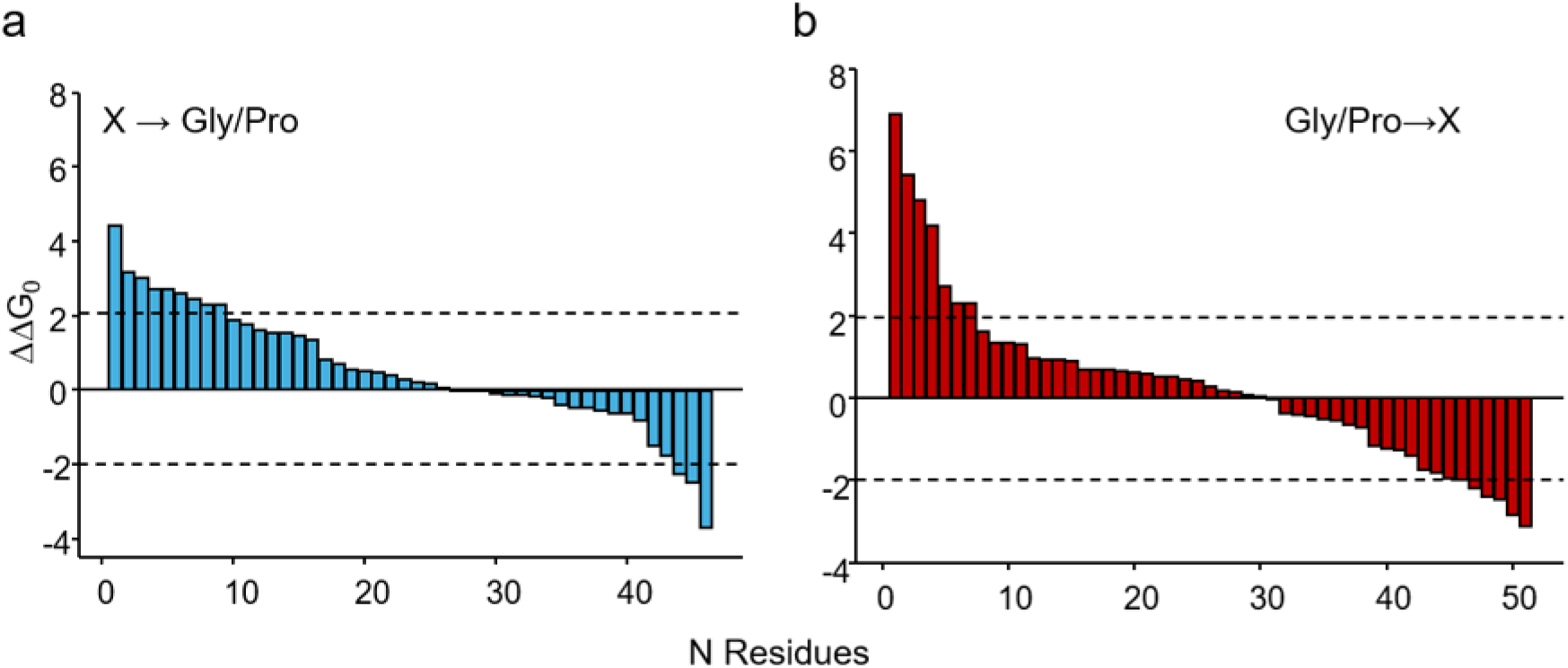
Backbone vs side-chain contributions to allosteric communication. Bar graphs show mutation-induced changes in ΔΔG_0_ for substitutions from glycine or proline to another amino acid, (X) **(b)**, and for reciprocal substitutions from (X) to glycine or proline **(a)**. The dotted lines indicate the ±2 kcal/mol range, within which the energetic effects of the two classes of substitutions are broadly comparable. Substantially larger free-energy changes are observed for substitutions of selected glycine and proline residues by other amino acids. These pronounced effects are restricted to strategically positioned glycine and proline residues within the receptor, particularly those located near the transmitter-binding sites, indicating an important contribution of local backbone conformational properties to AChR gating.

## DISCUSSION

### Parallel gating pathways reveal pre-existing allosteric routes in unliganded AChRs

Activation of allosteric receptors such as AChRs in the presence of agonists or mutations presumably involves 2 distinct steps: (i). initiation or trigger at the site of agonist binding or perturbation and (ii). communication of the information about allosteric signal initiation to the distant ion channel gate that regulates ion flow. The results presented here, including 1. dual gating kinetics of unliganded AChRs, with a predominant major- and a minor-(∼10%) long-opening component, 2. the presence of a distinct barrier between the two pathways, which is altered in the presence of some of the binding site mutations; lowered by some (e.g. αG153K) and elevated in a few other cases (e.g. αG147A), thereby either exaggerating or completely attenuating the long minor gating component, 3. the unliganded single channel data preferentially favouring a parallel-vs sequential-linear kinetic model, and 4. Distinct ϕ-values, and the relative stabilities of the gating vs long open-states, all indicate the presence of multiple parallel pathways. It has been proposed that the liganded neuromuscular AChRs has a broad, corrugated transition state landscape with multiple intermediate states separated by low-energy barriers (74). The different structural blocks of amino acids with distinct fractional ϕ-values were conceived by visualizing concerted, local conformational changes in the blocks in a sequential manner (conformational wave propagation (30)). However, observation of Hammond effect in AChRs (75) and the malleability of the TS in the presence of engineered mutations (68), strongly suggest that the TS landscape is amenable to change and alternative activation pathways are feasible.

Intrinsic gating energy (ΔΔG_0_) change and the expected gain of function (ΔΔG_2_^mut^) have been shown to be linearly correlated (with slope ≈ 1.0; see Fig. 3c) for a series of mutations at αA96 residue in the AChR (76). This affirms the idea that the major component of the unliganded gating, which has been classically utilized to study the kinetics and energetics of the unliganded AChRs (21, 47, 76), is a manifestation of the intrinsic gating free energy of the receptor. In contrast, the gating energy associated with the minor component of the unliganded gating approaches the ΔΔG_2_^ACh^ observed in the presence of ACh. The minor component is consistent with the earlier observations of modal gating in AChRs (24). Further, the ϕ-values of the minor component is significantly less than the major gating component and ≈ 0.1. Hence, in the reaction coordinate, this component seems to be ‘late’ compared to the major gating component. This makes sense as most of the time the receptor chooses the major pathway and only occasionally (probability ≈ 0.1) switches to the C_2_ state 0of the minor pathway. Low ϕ-values and preferential stabilization of the O state suggest that the receptor diffuses over a barrier-less TS once it reaches the minor C_2_- to O_L_ state.

### C-loop capping as the trigger for allosteric signal initiation

Loop-C has been shown to assume a capped, conformation, tightly wrapped around the agonists and uncapped conformation when bound to antagonists and toxins in AChBP structures, highlighting its importance in ligand binding (19). Recent cryo-EM and crystallographic studies on AChRs showed uncapped and capped conformational states of the C-loop in the Closed vs Open/Desensitized states (18, 63–65). Functionally, C-loop has been shown to be critical in determining agonist binding affinity (39, 62, 77) though constitutive gating was shown to be unaffected by the deletion of loop-C (62). Recently it was shown that in proton-activated chimeric Cys-loop receptors, deletion of all five loops C did not prevent channel opening or desensitization and suggested that the principal function of loop-C capping is local—promoting agonist capture and the low- to high-affinity reorganization of the transmitter-binding site (78). On the flip side, it was shown in simulations that forced closure of the C-loop leads to opening of the distant pore of the ion channel in AChRs (42). Further, mutation of C-loop residues to Gly leads to complete elimination of the minor gating component (62), though the major gating was intact. In this context, our results presented here including, 1. The modulation of the minor gating component by mutating the binding site and C-loop residues, 2. Occupancy of the C-loop in at least three distinct conformational states—uncapped, intermediate, and capped—even in the absence of ligand, 3. Strong correlation of the residence probability of the minor gating with the capped C-loop conformation and its close resemblance with the diliganded gating, suggest that loop C capping is critical in triggering the allosteric signal of an agonist occupying the binding pocket. In the unliganded receptor, the ∼10% occupancy in the long open state indicates that even in the absence of an agonist in the binding pocket, allosteric signal may be triggered every time the C-loop transitions to the capped conformation. The uncapped and intermediate capped conformations correspond to the predominant short-lived gating events. This establishes a direct structural–functional link between local binding-site conformational states and global channel gating behaviour.

In summary, the minor long-opening pathway is independent of agonist occupying the binding pocket. It is instead an intrinsic, low-probability route that becomes preferentially stabilized upon agonist binding. Thus, agonists do not impose a new pathway but rather select and amplify a pre-existing allosteric pathway.

### Non-cholinergic agonists reveal partial engagement of the allosteric network

The gating kinetics of non-cholinergic agonists (Fig. 1d) provides additional insight into the mechanism of signal initiation. These ligands produce complex single-channel kinetics resembling the unliganded receptor, suggesting incomplete or heterogeneous activation. Consistent with this, molecular dynamics simulations indicate that these ligands permit the binding pocket to sample all three C-loop conformations, rather than stabilizing the capped state in contrast with ACh. This implies that such agonists are inefficient at stabilizing the capped conformation, thereby failing to fully engage the high-efficacy pathway. These observations suggest that agonist efficacy is determined by the ability to stabilize C-loop capping, which in turn controls access to specific allosteric pathways. In this framework, partial agonists can be understood as ligands that do not effectively bias the conformational ensemble toward the capped state, resulting in reduced coupling between binding and gating.

### Allosteric communication differs fundamentally between unliganded and liganded receptors

Communications of allosteric signal from the binding site vs the sites of mutations in the receptor, intuitively ought to be different. In AChRs, our results suggest that allosteric signals in the presence of agonist is initiated by the capping of the C-loop, whereas in the unliganded receptor, local resettling triggers allosteric communication at the site of mutagenesis. Following initiation, the allosteric signal is communicated to the gate in nanoseconds time scale and is challenging to investigate. The current study highlights the fundamental difference in the allosteric communication between the unliganded (following mutagenesis) vs liganded receptors, as revealed by ϕ-value analysis and transition state studies. Although the overall energetic ranges are comparable, the distribution of ϕ-values differs substantially, indicating a reorganization of the TSE in the form of a conformational wave from the TBS to the gate (29, 30) in the presence of the agonist (Fig. 6) suggesting that agonist organizes the receptor into a state where allosteric communication is efficient and spatially continuous.

In contrast, the unliganded receptor lacks such a clearly defined pathway. Instead, the ϕ-values indicate that allosteric communication is more localized and originates from the site of perturbation (e.g., mutation site). This suggests that in the absence of ligand, the receptor exists in a loosely coupled network, where communication pathways are weakly coordinated.

#### Transition-state energetics and pathway selection

ΔΔG^‡^ estimates showed a coherent, highly coupled pathway linking the binding site to the channel gate, consistent with a “livewire”-like transmission of energetic perturbations in the liganded AChRs. The results imply that the location of the transition state along the reaction coordinate and the barrier heights determine allosteric communication. Allosteric communication involves lowering of the ΔΔG^‡^ values associated with these high-energy residues in the network. In contrast, in the unliganded receptor, the transition state is likely more distributed and less coordinated with few high-energy hub residues. This leads to a situation where multiple pathways compete, but none is strongly favoured, resulting in predominantly short-lived openings.

In summary, our results identify C-loop capping as the fundamental trigger for allosteric signal initiation in AChRs and reveal that ligand binding operates by selecting and stabilizing pre-existing allosteric pathways. This leads to the emergence of a highly efficient communication network linking the binding site to the channel gate. The default pathway for activation is mediated by C-loop capping in the presence of agonist. But, to prevent stray activation in the absence of agonist, in the unliganded receptor, this pathway has negligible probability. In constitutively active mutated AChRs, an altogether different pathway is chosen for activation, which perhaps has been perfected during evolution to attain maximum efficacy of the receptor.

## Acknowledgements

PK is grateful to CSIR for the research fellowship. The authors acknowledge Rachita Sharma for her insightful discussions and input to the project. The authors acknowledge IIT Delhi HPC facility for computational resources. The research was funded by SERB CRG (CRG/2022/007550), DBT (BT/PR47726/CMD/150/26/2023) and ICMR (IIRP-2023-0990) grant to TKN.

## Author contribution

Conceptualization: TKN; Execution of research, data analysis, and interpretation, review of manuscript: PK, and TKN; Original manuscript preparation: TKN and PK.

## Competing interest Statement

The authors declare no competing interests.

## Notes

### Competing Interest Statement

The authors have declared no competing interest.

