## Supplementary Tables for "Allosteric signal initiation and communication in neuromuscular acetylcholine receptors"

**Running Title:** Allosteric signaling in AChRs

**Authors**
Pradeepti Kampani[§], Tapan K. Nayak[§*]

**Author affiliations**
[§]Kusuma School of Biological Sciences, Indian Institute of Technology Delhi, Hauz Khas, New Delhi-110016

***Corresponding Author**

Tapan K. Nayak,
Kusuma School of Biological Sciences
Indian Institute of Technology Delhi,
Hauz Khas,
New Delhi-110016

**Author contribution**
Conceptualization: TKN; Execution of research, data analysis, and interpretation, review of manuscript: PK, and TKN; Original manuscript preparation: TKN and PK.

**Competing interest Statement:** The authors declare no competing interests.

**Key words:** C-loop capping, Activation barrier, Transition state, single-channel, patch-clamp, molecular simulations

**SI Table 1-** List of all kinetic models with their corresponding log likelihood scores used to determine the forward and backward rates for mutant constructs. Comparison of kinetic models based on LL scores for representative single channel data obtained from AChRs with background mutations at βL262S δL265S αD97A. Note the progressive increase in the LL scores with the number of free parameters. Model IX best described the single channel activity of this constitutive receptor.

| Construct | Model No | Model | No of free parameters | Log likelihood scores |
|---|---|---|---|---|
| αD97A βL262S δL265S | I | 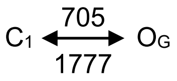 | 2 | 751842 |
| | II | 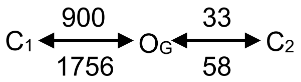 | 4 | 759909 |
| | III | 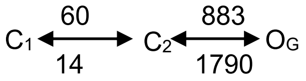 | 4 | 759909 |
| | IV | 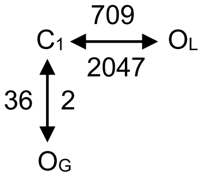 | 4 | 760469 |
| | V | 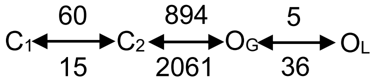 | 4 | 768564 |
| | VI | 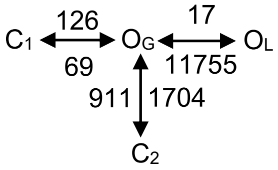 | 6 | 768564 |
| | VII | 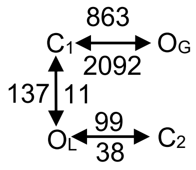 | 6 | 766343 |
| | VIII | 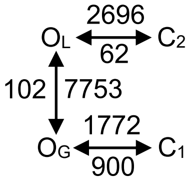 | 6 | 760221 |
| | IX | 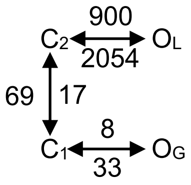 | 6 | 768864 |

| | | | | |
|---|---|---|---|---|
| | X | 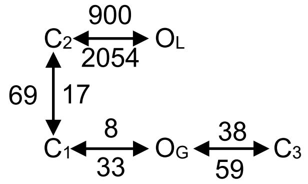  | 8 | 768532 |

**SI Table 2-** List of all kinetic models with their corresponding log likelihood scores used to determine the forward and backward rates for mutant constructs of and near the binding pocket. Comparison of kinetic models based on LL scores for representative single channel data obtained from AChRs with background mutations at αY93 αD97A αY127F αS269I. Model I best described the single channel activity of this constitutive receptor.

| **Construct** | **Model No** | **Model** | **No of free parameters** | **Log likelihood scores** |
|---|---|---|---|---|
| αY93 αD97A αY127F αS269I | I | 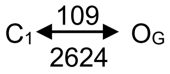  | 2 | 199482 |
| | II | 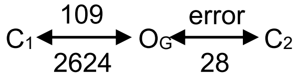  | 4 | 199461 |
| | III | 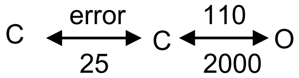  | 4 | 199462 |
| | IV | 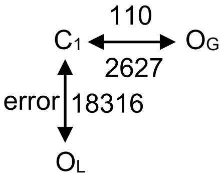  | 4 | 199450 |
| | V | 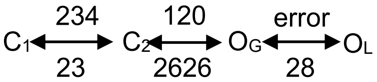  | 4 | 198823 |
| | VI | 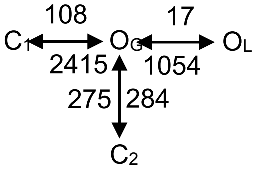  | 6 | 199468 |
| | VII | 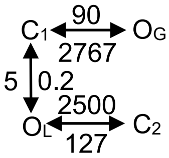  | 6 | 199741 |

| | | | | |
|---|---|---|---|---|
| | VIII | 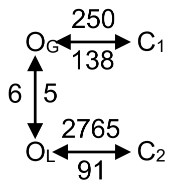 | 6 | 199740 |
| | IX | 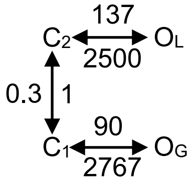 | 6 | 199741 |
| | X | 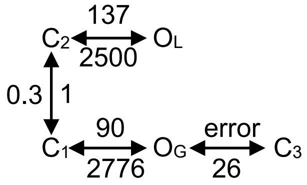 | 8 | 199745 |

**SI Table 3-** Kinetic rates and equilibrium constants for all mutant constructs used in the study

| **Background** | **Kinetic rate constants ($s^{-1}$)** | | | | | | | | | |
|---|---|---|---|---|---|---|---|---|---|---|
| | $C_1 \rightarrow O_G$ | $O_G \rightarrow C_1$ | $C_2 \rightarrow O_L$ | $O_L \rightarrow C_2$ | $C_1 \rightarrow C_2$ | $C_2 \rightarrow C_1$ | $C_1 \rightarrow O_3$ | $O_3 \rightarrow C_1$ | $C_2 \rightarrow O_G$ | $O_G \rightarrow C_2$ |
| αA96H<br>βT456I | 265 | 5031 | 2344 | 1021 | 104 | 2687 | - | - | - | - |
| αA96N<br>βT456I<br>αE181T<br>εL269F | 2380 | 2078 | 2465 | 652 | 1045 | 311 | - | - | - | - |
| αA96L<br>βL262Q<br>δL265Q | 5014 | 414 | 14214 | 126 | 725 | 63 | - | - | - | - |
| αA96Y<br>βL262F<br>δL265F | 254 | 26823 | 5840 | 6065 | 350 | 623 | - | - | - | - |
| αA96Y<br>δV269A | 4558 | 122 | 305 | 104 | 350 | 623 | - | - | - | - |
| αA96L<br>βL262Q<br>δL265Q | 6056 | 299 | 293 | 54 | 404 | 821 | - | - | - | - |
| αA96N<br>βV266A<br>δV269A | 3876 | 88 | 378 | 14380 | 419 | 452 | - | - | - | - |
| αA96Y<br>βV266Q<br>δV269q | 17529 | 74 | 707 | 9224 | 1469 | 2630 | - | - | - | - |
| αDYS<br>αG147A | 52 | 7138 | - | - | - | - | - | - | - | - |

| | | | | | | | | | | |
|---|---|---|---|---|---|---|---|---|---|---|
| αA96H αY93H | 202 | 7830 | - | - | - | - | - | - | 3296 | 2677 |
| αDYS αG147A αG153A | 272 | 6383 | | | | | | | | |
| αA96H εS450W polyG in loop C/ 3G | 534 | 1922 | - | - | - | - | - | - | 4454 | 773 |
| αA96N αW149S δV269A | 1304 | 58 | - | - | - | - | - | - | 12057 | 332 |
| αA96N αW149A βV266A | 2435 | 210 | - | - | - | - | - | - | - | - |
| αDYS αG153K | 2745 | 1752 | 664 | 175 | 55 | 299 | 600 | 519 | - | - |
| αDYS αP197A | | | | | | | | | | |
| αA96V αT456I εE181T εL269F | 209 | 13635 | 12235 | 2404 | 1191 | 0.26 | 0.6 | 236 | - | - |
| αA96L αT456I εE181W εL269F | 1740 | 5138 | 4233 | 1360 | 0.7 | 22.11 | 379 | 13.1 | - | - |
| αA96F βT456I εE181W εL269F | 2765 | 2578 | 2865 | 1015 | 13 | 155 | 67 | 285 | - | - |
| αD97A αY127F αS269I δN109Y | 356 | 191 | 194 | 5454 | 483 | 4.8 | 2471 | 3332 | - | - |

**SI Table 4.** Parallel gating pathway analysis in AChRs.

| **Residue** | **Mutation** | **Background** | **Closing rate from $O_G$ ($b_0$) ($s^{-1}$)** | **Closing rate from $O_L$ ($b_0'$) ($s^{-1}$)** | **$\Delta G_0$** | **$\Delta G_0^L$** | **$\Delta\Delta G_2$** |
|---|---|---|---|---|---|---|---|
| αA96 | C | βT456I δI43Q εE181T εL269F | 7634 | 3888 | - | - | - |
| αA96 | F | βT456I δI43Q εE181T εL269F | 1552 | 775 | -0.001 | -0.8 | -9.37 |
| αA96 | H | αY190R βT456I | 9217 | 4553 | | | |
| αA96 | H | αY93H | 15580 | 7107 | 0.068 | -0.55 | -6.06 |

| | | | | | | | |
|---|---|---|---|---|---|---|---|
| αA96 | N | δI43Q | 16663 | 7223 | 3.57 | 1.05 | -5.63 |
| αA96 | F | βT456I εE181W εL269F | 2578 | 1015 | -0.04 | -0.79 | -7.98 |
| αA96 | N | αW149S εE181T εL269F | 8788 | 3301 | - | - | - |
| αA96 | N | βT456I δI43Q εE181T εL269F | 6032 | 2264 | -0.45 | 1.1 | -9.75 |
| αA96 | N | αW149S βT456I δI43Q εE181T εL269F | 2093 | 785 | - | - | - |
| αA96 | L | βT456I εE181T εL269F | 10794 | 3478 | 1.674 | -0.12 | -6.14 |
| αA96 | L | βT456I εE181W εL269F | 5138 | 1360 | 0.63 | -0.66 | -7.09 |
| αA96 | L | βT456I δI43Q εE181W εL269F | 1448 | 3892 | 0.25 | -0.13 | -10.3 |
| αA96 | H | - | 10776 | 2489 | 2.52 | -0.37 | -6.88 |
| αA96 | H | αY93H αT254V δS268V | 1439 | 328 | 0.92 | -1.13 | -7.7 |
| αA96 | V | βT456I εE181T εL269F | 13635 | 2404 | 2.46 | -0.96 | -6.9 |
| αA96 | F | δI43Q εE181T εL269F | 2433 | 361 | 0.09 | -1.46 | -8.07 |
| αA96 | H | βT456I | 4618 | 509 | 1.45 | -0.53 | -7.3 |
| αA96 | E | βT456I εE181W εL269F | 4183 | 433 | 1.23 | -0.83 | -7.9 |
| αA96 | H | βT456I εE181T εL269F | 1309 | 130 | 0.51 | -1.5 | -10.72 |
| αA96 | N | εE181T εL269F | 5145 | 354 | 1.03 | -0.6 | -8.33 |
| αA96 | N | βT456I εE181T εL269F | 921 | 103 | -0.21 | -1.88 | -8.87 |
| αA96 | E | δI43Q εE181T εL269F | 1382 | 138 | -0.36 | -0.86 | -7.9 |
| αA96 | Y | δV269A | 1220 | 184 | -0.77 | -0.3 | -9.06 |
| αA96 | Y | βT456I εE181T εL269F | 1355 | 283.5 | -1.01 | -0.3 | -9.7 |
| αA96 | H | βT456I εE181T εL269F (150mM Na+) | 1222 | 176 | -1.02 | 0.94 | -10.73 |
| αA96 | H | βT456I εE181T εL269F (150mM Cs) | 858 | 70 | -1.4 | -2.07 | -10.73 |
| αA96 | H | βT456I εE181T εL269F (150mM K+) | 424 | 35 | -1.67 | -0.46 | -10.73 |

| | | | | | | | |
|---|---|---|---|---|---|---|---|
| αA96 | L | βL262Q δL265Q | 299 | 54 | -1.77 | -0.99 | -11.29 |
| αA96 | W | βT456I εE181T εL269F | 350 | 121 | -1.86 | -0.7 | -9.5 |
| αA96 | W | βT456I εE181W εL269F | 190 | 108 | -2.0 | -1.3 | -10.45 |
| αA96 | E | βT456I δI43Q εE181T εL269F | 558 | 1325 | - | - | - |
| αA96 0Cs+/dd H2O/ +70mV | Y | δV269A | 1125 | 384 | -0.56 | -0.57 | -9.06 |
| αA96 151Cs +/pipette /+70mV | Y | δV269A | 362 | 59 | -1.77 | -2.4 | -9.06 |
| βV266 | A | δV266A εE181W εL269F | 34 | 221 | -2.13 | -1.6 | -11.44 |
| αD97 | A | αY127F αS126F αW149S | 1189 | 553 | - | - | - |
| αD97 | A | αY127F αS126F βT265V δS268V | 413 | 648 | -1.28 | -0.45 | -7.7 |
| δN109 | Y | DYS | 5454 | 3332 | 1.97 | 0.17 | -6.56 |
| δW120 | F | DYS | 11334 | 5025 | 3.33 | 0.22 | -4.8 |
| δW120 | R | DYS | 2825 | 720 | 1.53 | -0.04 | -5.44 |
| εN182 | D | DYS | 3361 | 1617 | 1.06 | -0.93 | -5.74 |
| εN182 | D | αA96Y βT464I | 8308 | 2128 | 2.11 | -0.26 | -6.44 |
| αP197 | S | DYS | 2138 | 1456 | 0.92 | 0.65 | -6.97 |
| αP197 | A | DYS | 3301 | 1478 | 1.59 | -0.11 | -6.9 |
| αW149 | R | βT456I δI43Q εE181W εL269F | 3070 | 1170 | - | - | - |

Closing rate from $O_G$ ($b_0$), closing rate from $O_L$ ($b_0'$), $\Delta G_0$ (unliganded free energy change for major gating ($O_G$), $\Delta G_0{}^L$ (unliganded free energy change for major gating ($O_L$)

**SI Table 5**- Parallel gating pathway analysis in AChRs.

| **Residue** | **Mutation** | **Background** | **Closing rate from $O_G$ ($b_0$) ($s^{-1}$)** | **Closing rate from O'' ($b_0''$) ($s^{-1}$)** |
|---|---|---|---|---|
| αA96 | Y | εE181T εL269F | 3030 | 50 |
| αA96 | F | βT456I εE181W εL269F | 2578 | 171 |
| αA96 | H | βT456I εE181T εL269F | 1222 | 38 |

| | | | | |
|---|---|---|---|---|
| αA96 | H | βT456I εE181T εL269F (150mM Cs) | 858 | 26 |
| αA96 | H | βT456I εE181T εL269F (150mM K+) | 1424 | 35 |
| αA96 | L | βT456I εE181W εL269F | 5138 | 178 |
| αA96 | L | βT456I δ143Q εE181W εL269F | 14738 | 286 |
| αA96 | N | εE181T εL269F | 7145 | 224 |
| αA96 | V | βT456I εE181T εL269F | 13635 | 236 |
| αG153 | A | DYS | 1752 | 75 |
| αG153 | K | DYS | 779 | 55 |
| αP197 | D | DYS+G153K | 3057 | 99 |
| αP197 | A | DYS | 1448 | 80 |
| δN109 | Y | DYS | 5454 | 191 |
| δN109 | S | DYS | 2115 | 29 |

**SI Table 6**: $\Delta\Delta G_0$ for all residues containing side chains which are replaced with G (glycine) or P (proline).

| **X → Gly** | | **$\Delta\Delta G_0$ (kcal/mol)** |
|---|---|---|
| V46 | G | 4.4438 |
| Y93 | G | 3.1718 |
| 1260 | G | 3.0184 |
| Y127 | G | 2.7171 |
| V188 | G | 2.6095 |
| P265 | G | 2.3081 |
| P272 | G | 2.3081 |
| L42 | G | 1.8991 |
| V132 | G | 1.6218 |
| E45 | G | 1.5362 |
| δS258 | G | 1.3585 |
| K276 | G | 0.7103 |
| S275 | G | 0.5406 |
| εS256 | G | 0.5118 |
| I264 | G | 0.4911 |
| A96 | G | 0.4006 |
| δP286 | G | 0.2977 |
| εA271 | G | 0.2021 |
| S246 | G | 0.1938 |

| | | |
|---|---|---|
| L273 | G | 0.0556 |
| εP282 | G | 0.0344 |
| P120 | G | 7.12E-03 |
| R277 | G | -0.1076 |
| εT277 | G | -0.14 |
| M207 | G | -0.1557 |
| βE283 | G | -0.211 |
| F268 | G | -0.5638 |
| M415 | G | -0.6282 |
| E262 | G | -1.7675 |
| V249 | G | -2.2962 |
| S173 | G | -3.7038 |
| D97 | G1 | -0.8467 |
| D97 | G2 | -0.5091 |
| **X → Pro** | | **ΔΔ$G_0$ (kcal/mol)** |
| G275 | P | 2.7171 |
| Y127 | P | 2.4778 |
| αE45 | P | 1.7675 |
| Y127 | P | 1.5283 |
| L42 | P | 1.4902 |
| εL40 | P | 0.842 |
| δL40 | P | 0.0437 |
| S426 | P | -0.2069 |
| αA96 | P | -0.409 |
| S173 | P | -0.4845 |
| G153 | P | -2.49 |
| D97 | P1 | -0.6423 |
| D97 | P2 | 1.5078 |

**SI Table 7-** ΔΔ$G_0$ for G and P residues containing which are replaced with residues containing side chains.

| **Gly → X** | | **ΔΔ$G_0$ (kcal/mol)** |
|---|---|---|
| G147 | A | 5.44 |
| G147 | S | 4.82 |
| G147 | V | 6.91 |
| G153 | A | -2.41 |
| G153 | C | -2.87 |
| G153 | D | -2.0022 |
| G153 | K | -3.14 |

| | | |
|---|---|---|
| G153 | P | -2.49 |
| G153 | R | -1.8339 |
| G153 | S | -1.98 |
| G153 | W | -2.2167 |
| G153 | Y | -1.7769 |
| G174 | A | 0.4455 |
| G174 | D | 1.6406 |
| G174 | F | -0.394 |
| G174 | H | 0.0959 |
| G174 | R | 0.9798 |
| G174 | V | 0.1777 |
| G174 | W | -0.5335 |
| G174 | Y | 0.6194 |
| G275 | L | 0.6723 |
| G275 | P | 2.7171 |
| G275 | S | 0.9208 |
| eG183 | I | -0.6482 |
| eG183 | Y | -1.2831 |
| **Pro → X** | | **ΔΔ$G_0$ (kcal/mol)** |
| βP276 | A | -0.4652 |
| βP276 | D | -0.7391 |
| βP276 | H | 0.526 |
| βP276 | S | -0.4377 |
| δP279 | C | 0.9496 |
| δP279 | E | 0.9496 |
| δP279 | F | 1.3585 |
| δP279 | K | 1.3585 |
| δP279 | R | 0.7103 |
| δP279 | G | 0.2977 |
| εP275 | A | 0.15 |
| εP275 | E | 0.42 |
| εP275 | R | -0.57 |
| εP282 | G | 0.0344 |

**SI Table 8**- Values for $\phi_1$ (major component, $O_G$), $\phi_2$ (minor component, $O_L$), $\Delta G_{range}$ (kcal/mol) $\Delta\Delta G^‡$ (kcal/mol) for residues in α,β,δ,ε subunits of the adult unliganded AChR. The values are mapped onto the pdb structure PDB:9AWJ.

| **Human nAChR sequence** | **Phi1 ($\phi_1$)** | **$\Delta G_{range}$ (kcal/mol)** | **$\Delta\Delta G^‡$ (kcal/mol)** |
|---|---|---|---|
| **α subunit** | | | |
| E45 | 0.97 | -0.15421 | -0.1497 |

| | | | |
|---|---|---|---|
| I49 | - | -0.65 | 0 |
| T52 | 0.33 | -0.69575 | -0.23413 |
| D89 | 0.54 | -3.79784 | -2.05083 |
| Y93 | 0.83 | -1.98135 | -1.64452 |
| A96 | 0.71 | -6.92302 | -4.91535 |
| D97 | 0.31 | -3.089 | -0.95759 |
| S126 | 0.99 | -0.46 | -0.4554 |
| Y127 | 0.83 | -4.23048 | -3.5113 |
| G147 | 0.68 | -1.88627 | -1.28266 |
| I148 | 0.69 | -1.8027 | -1.24386 |
| W149 | 0.72 | -4.10353 | -2.95454 |
| T150 | 0.64 | -4.46223 | -2.85582 |
| Y151 | 0.66 | -2.62346 | -1.73148 |
| D152 | 0.56 | -0.13838 | 0 |
| G153 | 0.6 | -5.35832 | -3.21499 |
| F189 | 0.6 | -4.15144 | -2.49086 |
| Y190 | 0.89 | -0.48654 | -0.43302 |
| C192 | 0.84 | 0.189622 | 0.15928 |
| P197 | 0.61 | -2.57863 | -1.57296 |
| Y198 | 0.7 | -1.6138 | -1.12966 |
| R209 | 0.9 | -2.54 | -2.286 |
| N217 | 0.81 | -3.22924 | -2.61568 |
| V218 | 0.45 | -7.7 | -3.465 |
| P221 | 0.56 | -1.4 | -0.784 |
| F225 | 0.48 | -2.62 | -1.2576 |
| L251 | 0.39 | -6.14 | -2.3946 |
| T254 | 0.35 | -0.38 | -0.133 |
| V255 | 0.49 | -3.63 | -1.7787 |
| I260 | 0.82 | -5.36 | -4.3952 |
| E262 | 1 | -1.65 | -1.65 |
| P265 | 0.78 | -4.75 | -3.705 |
| S268 | 0.92 | -6.08 | -5.5936 |
| S269 | 0.46 | -2.28 | -1.05147 |
| P272 | 0.49 | -2.80385 | -1.37388 |
| L279 | 0.26 | -3.28472 | -0.85403 |
| I283 | 0.018 | -1.0459 | -0.01883 |
| C418 | 0.0421 | -3.72323 | -0.15675 |
| C193 | 0.83 | -0.59455 | -0.49347 |
| T422 | 0.15 | -1.966 | -0.2949 |
| **β subunit** | | | |
| N228 | - | -0.4 | 0 |
| V229 | - | -1.1 | 0 |
| P232 | - | -1.2 | 0 |
| I258 | 0.59 | -1.87 | -1.1033 |
| L262 | 0.43 | -4.88 | -2.0984 |
| T265 | 0.56 | -3.46093 | -1.93812 |

| | | | |
|---|---|---|---|
| V266 | 0.46 | -4.07 | -1.8722 |
| L269 | 0.58 | -1.04 | -0.6032 |
| **δ subunit** | | | |
| I43 | 0.76 | -1.70532 | -1.29604 |
| W57 | - | -1.35853 | 0 |
| N109 | 0.92 | -2.3703 | -2.18068 |
| W120 | 0.8 | -0.50687 | -0.4055 |
| L121 | 0.37 | -3.68344 | -1.36287 |
| D114 | 0.91 | -0.74511 | -0.67805 |
| S115 | 0.39 | -0.40181 | -0.15671 |
| L121 | 0.73 | -1.55263 | -1.13342 |
| P123 | 0.72 | -1.48596 | -1.06989 |
| P235 | - | -1 | 0 |
| N231 | - | -0.9 | 0 |
| L265 | 0.46 | -5.46 | -2.5116 |
| S268 | 0.56 | -4.2 | -2.352 |
| V269 | 0.55 | -3.5 | -1.925 |
| L272 | 0 | -0.44 | 0 |
| **ε subunit** | | | |
| K34 | 0.65 | -5.93543 | -3.85803 |
| W55 | 0.71 | -1.69828 | -1.20578 |
| E57 | - | -0.78 | 0 |
| L104 | 0.66 | -1.75646 | -1.15927 |
| L105 | 0.56 | -1.62714 | -0.9112 |
| C106 | 0.75 | -2.68311 | -2.01233 |
| N107 | 0.89 | -2.67389 | -2.37976 |
| P112 | 0.9 | -0.80676 | -0.72608 |
| D113 | 0.81 | -0.86697 | -0.70224 |
| Y117 | 0.28 | -1.81799 | -0.50904 |
| L119 | 1 | -0.58334 | -0.58334 |
| P121 | 0.68 | -1.43288 | -0.97436 |
| E181 | - | -0.45 | 0 |
| N182 | 0.89 | -1.04526 | -0.93028 |
| E184 | 0.76 | -0.86 | -0.6536 |
| N226 | - | -0.75 | 0 |
| P230 | - | -0.54 | 0 |
| L261 | 0.54 | -4.27 | -2.3058 |
| V265 | 0.57 | -5.57251 | -3.17633 |
| F268 | - | -1.3 | 0 |
| L269 | 0.55 | -3.48713 | -1.91792 |
| S450 | 0.34 | -1.5189 | -0.51643 |

**SI Table 9**- Values for $\phi_1$ (major component, $O_G$), $\Delta G_{range}$ (kcal/mol) $\Delta\Delta G^{\ddagger}$ (kcal/mol) for residues in α,β,δ,ε subunits of the adult liganded AChR. The values are mapped onto the pdb structure PDB:9AWJ.

| Human nAChR sequence | Phi (ϕ) | ΔGrange (kcal/mol) | ΔΔG‡ (kcal/mol) | Reference |
|---|---|---|---|---|
| α subunit | | | | |
| R20 | - | -0.7539 | - | (Mitra, 2005) |
| V29 | - | -0.2684 | - | (Mitra, 2005) |
| I31 | - | -0.4400 | - | (Mitra, 2005) |
| L35 | - | -0.2643 | - | (Mitra, 2005) |
| L40 | - | 8.4000 | - | (Mitra, 2005) |
| E45 | 0.8 | -6.5060 | -5.20481 | (Jha, 2012) |
| V46 | 0.81 | -5.0161 | -4.06307 | (Mitra, 2005) |
| N47 | 0.81 | -2.8280 | -2.29066 | (Mitra, 2005) |
| Q48 | 0.81 | -2.9652 | -2.40182 | (Mitra, 2005) |
| I49 | 0.71 | -2.8319 | -2.01068 | Prevost MS, 2012 |
| T32 | 1 | -1.1000 | -1.1 | |
| V54 | - | -1.9280 | 0 | (Mitra, 2005) |
| R55 | - | -0.5787 | 0 | (Mitra, 2005) |
| L56 | - | -0.3026 | 0 | (Mitra, 2005) |
| L65 | - | -0.1041 | 0 | (Mitra, 2005) |
| W86 | - | -2.0000 | 0 | Purohit, 2013 |
| P88 | - | 8.4000 | 0 | (Mitra, 2005) |
| D89 | - | -0.4075 | 0 | (Mitra, 2005) |
| Y93 | 0.88 | -4.6852 | -4.12294 | Purohit, 2012 |
| N94 | 0.86 | -3.8000 | -3.268 | Prevost MS, 2012 |
| A96 | 0.79 | -8.1834 | -6.46491 | Prevost MS, 2012 |
| D97 | 0.93 | -3.4960 | -3.25129 | Chakrapani, 2003 |
| L110 | - | -0.1481 | 0 | (Mitra, 2005) |

| | | | | |
|---|---|---|---|---|
| G114 | - | 8.4000 | 0 | (Mitra, 2005) |
| I116 | - | -0.0140 | 0 | (Mitra, 2005) |
| P120 | - | -0.0071 | 0 | (Mitra, 2005) |
| A122 | - | -0.7361 | 0 | (Mitra, 2005) |
| S126 | 0.79 | -3.3000 | -2.607 | Purohit, 2013 |
| Y127 | 0.77 | -7.4279 | -5.71947 | (Purohit, 2007) |
| V132 | 0.75 | -4.6162 | -3.46214 | (Jha,2007) |
| T133 | - | -1.1193 | 0 | (Jha,2007) |
| H134 | 0.71 | -1.3585 | -0.96455 | (Jha,2007) |
| F135 | 0.75 | -1.7053 | -1.27899 | (Jha,2007) |
| P136 | - | 0.0000 | 0 | (Jha,2007) |
| F137 | 0.78 | -0.9015 | -0.70319 | (Mitra, 2005) |
| D138 | 0.78 | -3.3863 | -2.64132 | (Mitra, 2005) |
| Q140 | 0.78 | -1.9294 | -1.50493 | (Mitra, 2005) |
| N141 | 0.78 | -0.8000 | -0.624 | (Mitra, 2005) |
| T143 | 0.78 | 0.1000 | 0.078 | (Mitra, 2005) |
| M144 | 0.84 | -1.9823 | -1.66515 | (Mitra, 2005) |
| K145 | 0.96 | -3.2577 | -3.12736 | (Purohit, 2007) |
| L146 | - | -0.0361 | 0 | (Mitra, 2005) |
| G147 | 0.86 | -0.3234 | -0.27812 | Purohit, 2011 |
| I148 | - | -0.4527 | 0 | |
| W149 | 0.97 | -5.5031 | -5.33798 | Purohit, 2012 |
| T150 | - | -1.0164 | 0 | |
| G153 | 0.96 | -3.1400 | -3.0144 | Purohit, 2011 |
| S173 | - | -3.7038 | 0 | |
| G174 | 0.82 | -2.1741 | -1.78273 | Cadugan, 2011 |
| E175 | 0.52 | -2.6963 | -1.4021 | Cadugan, 2011 |
| W176 | 0.85 | -1.2037 | -1.02317 | Cadugan, 2011 |
| V177 | - | -0.2930 | 0 | |

| | | | | |
|---|---|---|---|---|
| V188 | - | -2.6095 | 0 | |
| Y190 | 0.89 | -7.7768 | -6.92134 | Purohit, 2012 |
| Y198 | 1 | -5.8452 | -5.84516 | Purohit, 2012 |
| D200 | - | -4.4177 | 0 | |
| T202 | - | -0.1938 | 0 | |
| M207 | - | -2.4906 | 0 | |
| R209 | 0.72 | -3.6174 | -2.60455 | Jha, 2012 |
| L210 | - | -4.1490 | 0 | Jha, 2012 |
| P211 | - | -0.7103 | 0 | Jha, 2012 |
| L212 | - | -1.2000 | 0 | Purohit, 2013 |
| Y213 | 0.8 | -2.4000 | -1.92 | Purohit, 2013 |
| F214 | 0.32 | -3.2000 | -1.024 | Purohit, 2013 |
| V215 | - | -1.0000 | 0 | Purohit, 2013 |
| V216 | - | -0.7000 | 0 | Purohit, 2013 |
| N217 | 0.79 | -3.1000 | -2.449 | Purohit, 2013 |
| V218 | 0.59 | -8.4000 | -4.956 | Purohit, 2013 |
| I219 | - | -1.2323 | 0 | Purohit, 2013 |
| I220 | - | -0.6000 | 0 | Purohit, 2013 |
| P221 | - | -4.9000 | 0 | Purohit, 2013 |
| C222 | 0.58 | -4.3000 | -2.494 | Purohit, 2013 |
| L223 | - | -1.4000 | 0 | Purohit, 2013 |
| L224 | 0.57 | -1.8000 | -1.026 | Purohit, 2013 |
| F225 | 0.57 | -3.3000 | -1.881 | Purohit, 2013 |
| S226 | 0.59 | -4.0000 | -2.36 | Purohit, 2013 |
| F227 | 0.58 | -1.5315 | -0.88825 | Purohit, 2013 |
| L228 | 0.67 | -3.0000 | -2.01 | Purohit, 2013 |
| T229 | - | -0.5702 | 0 | Purohit, 2013 |
| S220 | - | -1.2187 | 0 | Purohit, 2013 |

| | | | | |
|---|---|---|---|---|
| L231 | 0.59 | -1.8000 | -1.062 | Purohit, 2013 |
| V232 | - | -0.2194 | 0 | Purohit, 2013 |
| F233 | - | -1.2752 | 0 | Purohit, 2013 |
| Y234 | - | -0.8000 | 0 | Purohit, 2013 |
| L235 | - | -1.3000 | 0 | Purohit, 2013 |
| P236 | - | NF | 0 | Purohit, 2013 |
| T237 | - | -1.1000 | 0 | Purohit, 2013 |
| D238 | - | -0.5010 | 0 | Purohit, 2013 |
| S239 | - | -0.9528 | 0 | Purohit, 2013 |
| G240 | - | -0.3000 | 0 | Purohit, 2013 |
| T244 | - | 0.0000 | 0 | Purohit, 2007 |
| L245 | 0.61 | -2.0182 | -1.23108 | Purohit, 2007 |
| S246 | 0.63 | -2.9156 | -1.83681 | Purohit, 2007 |
| I247 | - | -2.5000 | 0 | Purohit, 2007 |
| S248 | 0.65 | -1.2037 | -0.78242 | Purohit, 2007 |
| V249 | 0.5 | -2.4719 | -1.23595 | Purohit, 2007 |
| L250 | 0.66 | -1.3023 | -0.85951 | Purohit, 2007 |
| L251 | 0.26 | -4.5600 | -1.1856 | Purohit, 2007 |
| S252 | - | 0.0000 | 0 | Purohit, 2007 |
| L253 | 0.55 | -3.1588 | -1.73734 | Purohit, 2007 |
| T254 | 0.35 | -3.8543 | -1.34899 | Purohit, 2007 |
| V255 | 0.52 | -5.5018 | -2.86096 | Purohit, 2007 |
| F256 | 0.67 | -2.9381 | -1.96854 | Purohit, 2007 |
| L257 | 0.62 | -3.7132 | -2.30221 | Purohit, 2007 |

| | | | | |
|---|---|---|---|---|
| L258 | 0.61 | -3.3044 | -2.01567 | Purohit, 2007 |
| V259 | 0.61 | 0.0000 | 0 | Mitra, 2005 |
| I260 | 0.89 | -3.4274 | -3.05038 | Bafna,2008 |
| V261 | 0.78 | -4.0756 | -3.17895 | Bafna,2008 |
| E262 | 0.82 | -3.0806 | -2.52609 | Bafna,2008 |
| L263 | 0.66 | -3.2578 | -2.15012 | Bafna,2008 |
| I264 | 0.78 | -4.6271 | -3.60912 | Bafna,2008 |
| P265 | 0.9 | -3.5495 | -3.19458 | Bafna,2008 |
| S266 | 0.64 | -4.0756 | -2.60837 | Bafna,2008 |
| T267 | 0.71 | -2.8487 | -2.02258 | Bafna,2008 |
| S268 | 0.97 | -5.0100 | -4.85969 | Bafna,2008 |
| S269 | 0.61 | -1.3984 | -0.85305 | Mitra, 2005 |
| A270 | 0.65 | -2.9548 | -1.92063 | Jha, 2007 |
| V271 | - | -0.4056 | 0 | Jha, 2007 |
| P272 | 0.62 | -7.0625 | -4.37875 | Jha, 2007 |
| L273 | - | -1.7330 | 0 | Jha, 2007 |
| I274 | 0.62 | -4.4845 | -2.78041 | Jha, 2007 |
| G275 | 0.65 | -2.7171 | -1.76608 | Jha, 2007 |
| K276 | - | -0.9496 | 0 | Jha, 2007 |
| Y277 | 0.34 | -2.0498 | -0.69693 | Cadugan, 2007 |
| M278 | - | 1.2000 | 0 | |
| L279 | 0.27 | -2.9810 | -0.80486 | Cadugan, 2007 |
| F280 | 0.3 | -0.8000 | -0.24 | Cadugan, 2007 |
| T281 | 0.63 | -2.5000 | -1.575 | Purohit, 2013 |
| M282 | - | -0.6000 | 0 | Purohit, 2013 |
| I283 | - | -1.3957 | 0 | Cadugan, 2007 |
| F284 | 0.32 | -1.9000 | -0.608 | Purohit, 2013 |
| V285 | - | 0.0000 | 0 | Purohit, 2013 |
| I286 | - | 0.0000 | 0 | Purohit, 2013 |
| A287 | - | 0.0000 | 0 | Purohit, 2013 |
| S288 | 0.61 | -2.4000 | -1.464 | Purohit, 2013 |
| I289 | 0.53 | -2.9000 | -1.537 | Purohit, 2013 |
| I290 | 0.31 | -1.0571 | -0.32771 | Cadugan, 2007 |

| | | | | |
|---|---|---|---|---|
| I311 | - | 0.0000 | 0 | Purohit, 2013 |
| T292 | 0.48 | -1.0000 | -0.48 | Purohit, 2013 |
| V293 | - | -0.1400 | - | Cadugan, 2007 |
| I294 | - | -0.2900 | - | Cadugan, 2007 |
| V295 | - | 0.0000 | - | Purohit, 2013 |
| I296 | - | -0.8000 | - | Purohit, 2013 |
| N297 | - | -0.2400 | - | Cadugan, 2007 |
| T298 | - | 0.0000 | - | Purohit, 2013 |
| H299 | - | -0.2000 | - | Purohit, 2013 |
| L410 | 0.54 | -0.9950 | -0.53729 | Mitra, 2004 |
| V413 | - | 0.0000 | - | Purohit, 2013 |
| M415 | 0.57 | -0.9296 | -0.52985 | Mitra, 2004 |
| V417 | - | 0.0000 | - | Purohit, 2013 |
| C418 | 0.5 | -3.2085 | -1.60423 | Mitra, 2004 |
| I420 | - | 0.0000 | - | Purohit, 2013 |
| T422 | 0.54 | -1.5000 | -0.81 | Mitra, 2004 |
| L423 | - | 0.0000 | - | Purohit, 2013 |
| A424 | - | 0.0000 | - | Purohit, 2013 |
| F426 | 0.56 | -2.2536 | -1.26201 | Mitra, 2004 |
| A427 | - | 0.0000 | - | Purohit, 2013 |
| G428 | - | 0.0000 | - | Purohit, 2013 |
| R429 | - | 0.0000 | - | Purohit, 2013 |
| S434 | - | -1.3817 | - | Purohit, 2013 |
| **β subunit** | | | | |
| D96 | - | -0.4221 | - | Purohit, 2013 |
| G97 | - | -0.6451 | - | Purohit, 2013 |
| F134 | - | -0.1985 | - | Purohit, 2013 |

| | | | | |
|---|---|---|---|---|
| D137 | - | -1.0529 | - | Purohit, 2013 |
| K220 | - | -0.0207 | - | Purohit, 2013 |
| E282 | - | -0.2110 | - | Purohit, 2013 |
| I283 | - | -0.4999 | - | Purohit, 2013 |
| L279 | - | -1.5570 | - | Purohit, 2013 |
| S278 | 0.17 | -2.4408 | -0.41493 | Purohit, 2013 |
| T277 | - | 0.0000 | - | Purohit, 2013 |
| E276 | 0.4 | -1.6268 | -0.65073 | Purohit, 2013 |
| P275 | - | -1.2652 | - | Purohit, 2013 |
| V274 | - | -1.0993 | - | Purohit, 2013 |
| K273 | 0.14 | -1.2964 | -0.18149 | Purohit, 2013 |
| D272 | - | -0.2450 | - | Purohit, 2013 |
| A271 | - | -0.5325 | - | Purohit, 2013 |
| L270 | - | -1.0455 | - | Purohit, 2013 |
| L269 | 0.4 | -6.3128 | -2.52512 | Purohit, 2013 |
| L268 | 0.63 | -2.8581 | -1.80058 | Purohit, 2013 |
| L267 | - | -1.2510 | - | Purohit, 2013 |
| F266 | - | -0.2916 | - | Purohit, 2013 |
| V265 | 0.13 | -5.8893 | -0.76561 | Purohit, 2013 |
| T264 | 0.39 | -2.6000 | -1.014 | Purohit, 2013 |
| L263 | - | 0.0000 | - | Purohit, 2013 |
| T262 | 0 | -1.6171 | 0 | Purohit, 2013 |
| L261 | 0.3 | -5.8000 | -1.74 | Purohit, 2013 |
| L260 | 0.32 | -2.6212 | -0.83877 | Purohit, 2013 |

| | | | | |
|---|---|---|---|---|
| A259 | 0.44 | -1.8701 | -0.82285 | Purohit, 2013 |
| F258 | - | -1.3826 | - | Purohit, 2013 |
| I257 | - | -2.0733 | - | Purohit, 2013 |
| S256 | - | -0.7019 | - | Purohit, 2013 |
| L255 | - | -1.3585 | - | Purohit, 2013 |
| G254 | - | 0.0000 | - | Purohit, 2013 |
| M253 | - | -1.6599 | - | |
| P231 | - | -2.0000 | - | |
| V228 | - | -2.5800 | - | |
| N227 | - | -0.4000 | - | |
| T464 | 0.17 | - | - | Mitra, 2004 |
| **δ subunit** | | | | |
| M283 | - | -0.7666 | - | Purohit, 2013 |
| S258 | - | -2.2327 | - | Purohit, 2013 |
| I261 | -0.05 | -1.4148 | 0.070738 | Purohit, 2013 |
| S262 | - | -0.7391 | - | Purohit, 2013 |
| S282 | - | -0.6863 | - | Purohit, 2013 |
| T281 | - | -1.3585 | - | Purohit, 2013 |
| A280 | - | -0.7103 | - | Purohit, 2013 |
| P279 | 0.56 | -1.3585 | -0.76077 | Purohit, 2013 |
| L278 | - | -0.9496 | - | Purohit, 2013 |
| R277 | 0.59 | -1.4661 | -0.865 | Purohit, 2013 |
| K276 | - | -1.4410 | - | Purohit, 2013 |
| S275 | 0.31 | -2.8246 | -0.87563 | Akk, 2013 |
| I274 | - | -1.5133 | - | Purohit, 2013 |
| L273 | 0.37 | -2.9000 | -1.073 | Akk, 2013 |
| L272 | 0.26 | -3.9700 | -1.03219 | Purohit, 2013 |
| L271 | - | -1.0000 | - | Purohit, 2013 |

| | | | | |
|---|---|---|---|---|
| L278 | - | -0.9496 | - | Purohit, 2013 |
| F270 | - | -1.0000 | - | Purohit, 2013 |
| L265 | 0.11 | -4.1487 | -0.45636 | Akk, 2013 |
| V269 | 0.44 | -3.4100 | -1.5004 | Purohit, 2013 |
| S268 | 0.28 | -4.0806 | -1.14257 | Purohit, 2013 |
| Q267 | - | -1.1000 | - | Purohit, 2013 |
| A266 | -0.05 | -1.2000 | 0.06 | Purohit, 2013 |
| L265 | 0.11 | -4.3000 | -0.473 | Akk, 2013 |
| L264 | - | -1.0000 | - | Purohit, 2013 |
| V263 | -0.1 | -1.0000 | 0.1 | Akk, 2013 |
| S262 | - | -0.6000 | - | Purohit, 2013 |
| I261 | -0.05 | -0.7000 | 0.035 | Akk, 2013 |
| A260 | - | -0.8000 | - | Purohit, 2013 |
| V259 | - | -1.0000 | - | Purohit, 2013 |
| S258 | - | -1.0952 | 2.1 | Akk, 2013 |
| T257 | - | 0.0000 | - | - |
| E45 | - | -1.5255 | - | - |
| K46 | - | -0.0909 | - | - |
| I43 | 0.86 | -1.8991 | -1.63326 | Purohit, 2007 |
| W57 | 0.94 | -1.6681 | -1.56803 | Bafna, 2009 |
| P123 | 0.9 | -5.1000 | -4.59 | Gupta, 2013 |
| L40 | - | -0.0437 | - | |
| D140 | - | -0.1697 | - | |
| F137 | - | -1.4683 | - | |
| F139 | - | -0.5188 | - | |
| K224 | - | -0.3058 | - | |
| L287 | - | -0.3793 | - | |
| L42 | - | -1.8991 | - | |
| N41 | - | -4.1044 | - | |
| E47 | - | -0.4202 | - | |
| V48 | - | -0.0325 | - | |
| M283 | - | -0.0732 | - | |
| P286 | - | -0.2977 | - | |
| **ε subunit** | | | | |
| S278 | - | -0.8800 | 0.8 | Jha, 2009 |

| | | | | |
|---|---|---|---|---|
| T277 | - | -0.8300 | - | Jha, 2009 |
| E276 | 0.56 | -1.5600 | -0.8736 | Jha, 2009 |
| P275 | 0.51 | -0.9900 | -0.5049 | Jha, 2009 |
| I274 | - | -1.2000 | - | Jha, 2009 |
| K273 | - | -1.0800 | - | Jha, 2009 |
| Q272 | - | -1.2100 | - | Jha, 2009 |
| A271 | - | -1.2929 | - | Jha, 2009 |
| I270 | - | -1.0455 | - | Jha, 2009 |
| L269 | 0.52 | -3.0606 | -1.59149 | Jha, 2009 |
| F268 | 0.51 | -3.7709 | -1.92316 | Jha, 2009 |
| L267 | - | -1.2037 | - | Jha, 2009 |
| F266 | 0.56 | -2.9612 | -1.65826 | Jha, 2009 |
| V265 | 0.34 | -5.6934 | -1.93577 | Jha, 2009 |
| T264 | 0.26 | -5.2872 | -1.37467 | Jha, 2009 |
| Q263 | 0.55 | -1.9192 | -1.05555 | Jha, 2009 |
| A262 | 0.59 | -1.7020 | -1.0042 | Jha, 2009 |
| L261 | 0.37 | -4.1989 | -1.5536 | Jha, 2009 |
| L260 | 0.5 | -2.8368 | -1.41839 | Jha, 2009 |
| V259 | 0.58 | -1.6901 | -0.98026 | Jha, 2009 |
| N258 | - | -1.1600 | - | Jha, 2009 |
| I257 | 0.33 | -5.5782 | -1.84081 | Jha, 2009 |
| S256 | - | -1.0812 | - | Jha, 2009 |
| V255 | 0.56 | -2.0718 | -1.16021 | Jha, 2009 |
| T254 | - | -0.6723 | - | Jha, 2009 |
| C253 | - | -0.4000 | - | |
| W55 | 0.97 | -1.6716 | -1.62145 | Bafna, 2009 |
| P121 | 0.98 | -5.2000 | -5.096 | Gupta, 2013 |
| E181 | 0.89 | -2.4599 | -2.18927 | Cadugan, 2010 |
| N182 | 0.78 | -2.0683 | -1.61326 | Cadugan, 2010 |
| G183 | 0.97 | -1.2831 | -1.24461 | Cadugan, 2010 |
| E184 | 0.8 | -3.2577 | -2.60613 | Cadugan, 2010 |
| S450 | 0.33 | -1.6800 | -0.5544 | Mitra, 2004 |

**SI Table 10**- Structural analysis of data obtained from MD simulation to understand loop C- and binding site dynamics for the WT vs mutant AChRs.

| **Residue** | **Mutation** | **Distance between C192 (loop C) and W149 (loop B) (Å) (mean ± S.D.)** | | | **Binding pocket volume (Å$^3$) (mean ± S.D.)** | |
|---|---|---|---|---|---|---|
| | | *Uncapped* | *Intermediate* | *Capped* | *Uncapped* | *Capped* |
| WT | - | 20.9±1.02 | 12.8±1.09 | 5±1.07 | 260 | 140 |

| | | | | | | |
|---|---|---|---|---|---|---|
| G147 | A | 24.0±0.96 | 15.2±0.86 | - | 250 | - |
| G153 | K | 22.9±0.96 | 10±1.2 | 4.8±0.60 | 300 | 130 |
| P197 | A | - | 6.3±1.3 | 3.2±1 | - | 120 |
| eE181 | W | 21.02±0.56 | 16.51±0.69 | - | 389, 302 | - |

**SI Table 11**- Structural analysis of data obtained from MD simulation to understand loop C- and binding site dynamics of the WT AChRs in the presence of cholinergic and non-cholinergic agonists.

| **Ligand** | **Distance between C192 (loop C) and W149 (loop B) (Å) (mean ± S.D.)** | | | **Binding pocket volume (Å$^3$) (mean ± S.D.)** | |
|---|---|---|---|---|---|
| | *Uncapped* | *Intermediate* | *Capped* | *Uncapped* | *Capped* |
| ACh | - | 9±1.2 | 6±1 | - | 160 |
| Betaine | - | 6±0.83 | 2.2±0.82 | - | 130 |
| Nicotinamide | - | 11±0.23 | 4.89±2.3 | - | 150 |

**SI Table 12**- State residence probabilities obtained from the MD simulations of WT and mutant AChRs and single channel state residence probabilities obtained from mean open time constants and the area under the major and minor components in dwell-time duration distribution histograms.

| **Residue** | **Mutation** | **Residence probability (*Simulation*)** | | | **Residence probability (*Experiment*)** | | |
|---|---|---|---|---|---|---|---|
| | | *Uncapped* | *Intermediate* | *Capped* | *Uncapped* | *Intermediate* | *Capped* |
| Unliganded WT | - | 0.7207 | - | 0.2793 | 0.4464 | - | 0.5536 |
| G147 | A | - | - | 1 | - | - | 0.9672 |
| G153 | K | 0.5968 | 0.0410 | 0.3622 | 0.6463 | - | 0.3389 |
